# Structure of the turnip yellows virus particles

**DOI:** 10.1101/2024.10.14.617924

**Authors:** Joséphine Lai-Kee-Him, Stefano Trapani, Sylvaine Boissinot, Catherine Reinbold, Chloé Fallet, Aurélie Ancelin, François Lecorre, François Hoh, Véronique Ziegler-Graff, Véronique Brault, Patrick Bron

**Affiliations:** Centre de Biologie Structurale (CBS), Univ Montpellier, CNRS, INSERM, Montpellier, France; INRAE, Université de Strasbourg, UMR SVQV, Colmar France; Institut de biologie moléculaire des plantes, CNRS, Université de Strasbourg, Strasbourg, France

**Keywords:** polerovirus, readthrough domain, capsid, elevated dodecahedron, structure, electron cryo-microscopy, immunogold labelling

## Abstract

Turnip yellows virus (TuYV) is a plant virus infecting important crops such as oilseed rape. TuYV is phloem-restricted and transmitted by aphids. The capsid contains two subunit types: the major capsid protein (CP) and a minor component (RTP*) which arises from the C-terminal cleavage of a readthrough product (RTP). RTP* contains the CP sequence fused with a structured domain, denoted ^N^RTD, which is a key determinant of virus transmission. Though both CP and RTP* are involved in virus movement and aphid transmission, how RTP* is incorporated into the capsid is poorly understood. We present here the structural characterisation, by immunogold labelling and 3D cryo-EM, of the wild-type TuYV and a mutant whose capsid contains the CP only. We show that incorporation of RTP* does not impair the capsid structure, and the ^N^RTD does not adopt well-defined positions at the capsid surface. The number of incorporated RTP*s suggests a random insertion.

## Introduction

Turnip yellows virus (TuYV), formerly beet western yellows virus, is a member of the genus *Polerovirus*. Members of the genera *Polerovirus* and *Enamovirus* (family *Solemoviridae*) and *Luteovirus* (family *Tombusviridae*), hereafter referred to as P/E/L viruses (Schiltz et al., 2022), are important plant pathogens that affect many major crops (Heck and Brault, 2018). Their economic importance extends worldwide to small grains, potato, brassicas, cotton, cucurbits, pepper, sugarcane and sorghum. P/E/L viruses are phloem-restricted and transmitted exclusively by aphids in nature (Gildow, 1999). The P/E/L genome is a linear single-stranded positive-sense RNA with an intricate organisation of the open reading frames (ORFs) (Boissinot et al., 2020; Smirnova et al., 2015; Sõmera et al., 2021) (Fig. 1a). The genome is encapsidated into a non-enveloped particle of ∼30 nm in diameter composed of 180 subunits arranged according to a *T*=3 icosahedral symmetry. Most of the 180 subunits are the major capsid protein (CP) of about 22-25 kDa encoded by ORF3. However, some of the subunits (10 to 25%) contain a C-terminal extension which protrudes at the virion surface (Brault et al., 1995; Filichkin et al., 1994). This minor capsid protein, hereafter called RTP* (∼55 kDa), arises from stochastic translational readthrough of the CP stop codon, whereby a non-structural readthrough protein (RTP) of about 74 kDa is produced (Fig. 1b). The RTP contains the CP sequence on the N-ter side, followed by a proline-rich flexible linker and a readthrough domain (RTD). This is composed of a structured sub-domain, denoted ^N^RTD (Schiltz et al., 2022), and a C-ter sub-domain, denoted ^C^RTD, which is predicted to be intrinsically disordered (LaTourrette et al., 2021). Post-translational cleavage of the ^C^RTD stretch from the RTP results in the structural RTP* which is incorporated in the capsids (Fig. 1b).

**Fig. 1.**
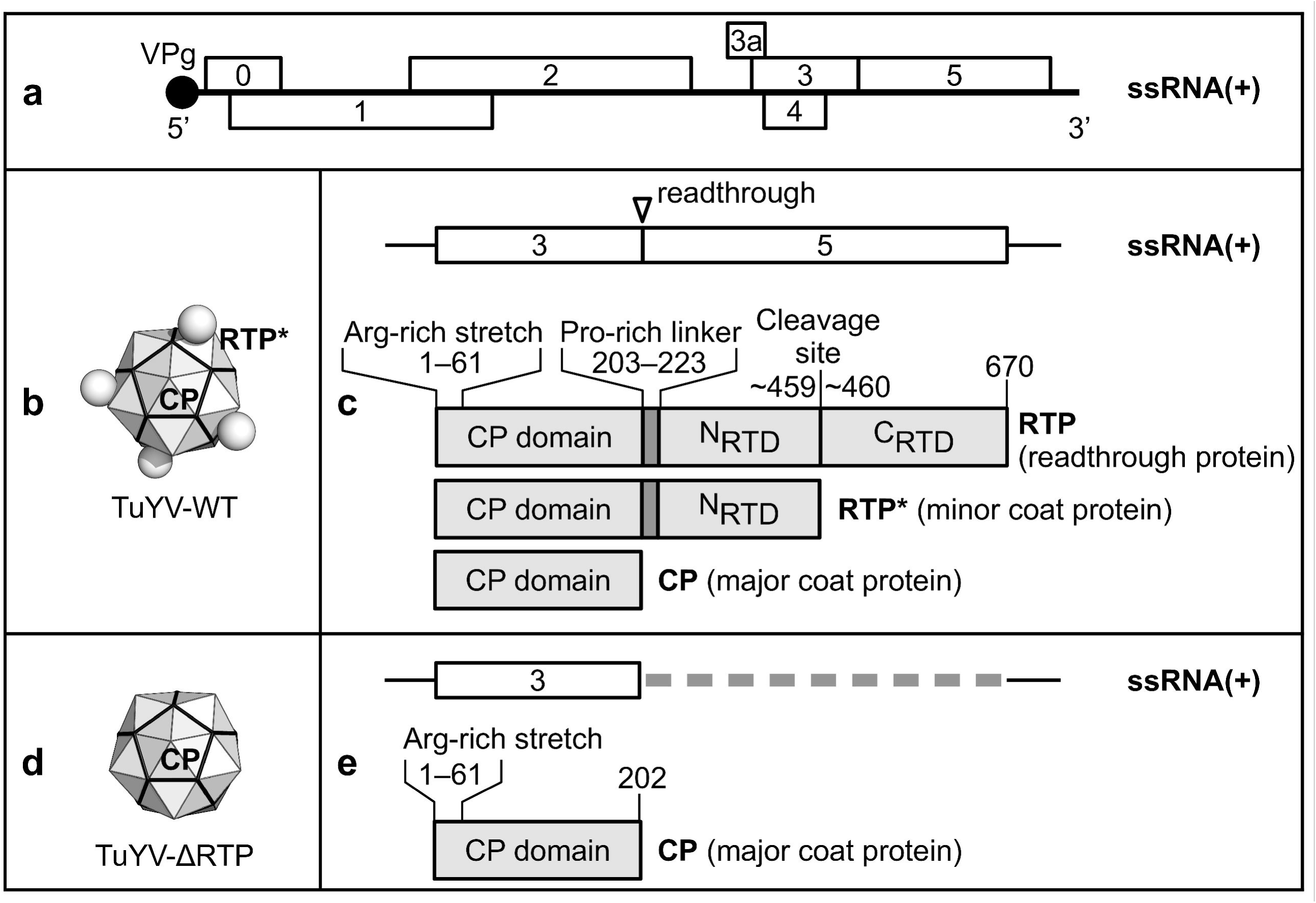
Encoding and organisation of the structural proteins in TuYV. (**a**) TuYV genome organisation (5641 nucleotides). The numbered rectangles represent ORFs, whose functions in the virus cycle are described in Boissinot et al. (2020). The VPg capping is indicated at the genome 5’ end. (**b**) Schematic representation of a wild-type virion (TuYV-WT). The capsid is composed of a mixture of the major coat protein (CP) and the minor coat protein (RTP*). An (arbitrary) number of ^N^RTD extensions (from the RTP*) are represented by small spheres. (**c**) Representation of the ORF3 and 5 in the wild-type virus and their expression products (RTP, RTP* and CP). The readthrough position at the ORF3 stop codon is indicated by a triangle. The amino-acid numbers delimiting the arginine-rich N-terminal stretch, the proline-rich linker, and the putative cleavage site on the RTP are shown. (**d**) Schematic representation of the TuYV-ΔRTP mutant, whose capsid is composed exclusively of the major coat protein. (**e**) as in (**c**) for the TuYV-ΔRTP mutant. The deletion of the ORF5 sequence is indicated by a dotted line.

The role of RTP, RTP* and CP in P/E/L virus movement and transmission by aphids has been extensively addressed, based mainly on reverse genetics on TuYV, potato leafroll virus (PLRV, genus *Polerovirus*), barley yellow dwarf virus (BYDV, genera *Luteovirus* and *Polerovirus* depending on the species) and cucurbit aphid-borne yellows virus (CABYV, genus *Polerovirus*). The full-length non-structural RTP is involved in virus systemic invasion of the plant (Boissinot et al., 2014; Brault et al., 1995; Hipper et al., 2014; Kaplan et al., 2007; Lee et al., 2005; Mutterer et al., 1999; Peter et al., 2008; Rodriguez-Medina et al., 2015; Wang et al., 1995; Xu et al., 2018) and the terminal ^C^RTD plays a role in restricting PLRV to the phloem (Peter et al., 2009) and in symptom development (Bruyère et al., 1997; Rodriguez-Medina et al., 2015). The RTP* is not required for capsid assembly (Bruyère et al., 1997; Reutenauer et al., 1993), but the presence of both CP and RTP* in intact virions is required for the virus to move long distance within host plants and for transmission by aphid vectors (Brault et al., 1995; Chay et al., 1996; Hipper et al., 2014; Lee et al., 2005). It has been shown that both TuYV structural proteins interact with the ephrin receptor, an aphid protein involved in virus uptake and transmission (Mulot et al., 2018). Also, the ^N^RTD of RTP* has been identified as a key determinant of transmission as mutant viruses devoid of the whole RTD are not aphid transmissible and retain the ability to infect plants only at a reduced titre (Brault et al., 1995; Chay et al., 1996; Liang et al., 2004; Reutenauer et al., 1993). Conversely, mutants lacking just the ^C^RTD assemble efficiently into virions (Boissinot et al., 2014; Bruyère et al., 1997), retain the ability to interact with aphid proteins (Mulot et al., 2018) and can be transmitted to new hosts (Wang et al., 1995). *In vitro* and *in vivo* binding assays also revealed that TuYV and PLRV virions bind to plant components (Bencharki et al., 2010; DeBlasio et al., 2016, 2015a, 2015b).

The structure of P/E/L capsids and how they interact with aphid vectors have been the object of investigation for several years. Difficulties in producing enough purified viruses had long limited structural studies to homology modelling coupled to mutational and cross-linking experiments (Alexander et al., 2017; Brault et al., 2003; Chavez et al., 2012; Kaplan et al., 2007; Terradot et al., 2001; Torrance, 1992). Significant advances were made after the production of BYDV and PLRV virus-like particles (VLPs) containing the recombinant CP, but not the RTP*, and the establishment of their atomic structure by electron cryogenic microscopy (cryo-EM) (Byrne et al., 2019). Also, the crystal structure of the ^N^RTD, absent from the VLPs, was solved using recombinant forms of the isolated domains from PLRV and TuYV (Schiltz et al., 2022). These studies revealed that P/E/L viruses are structurally related to picorna-like viruses, corrected errors in previously proposed structural models, and rationalised the observations from previous mutagenesis studies. The ^N^RTD structure revealed a dimeric architecture in which the sequence of each monomer goes back and forth from a jell-roll domain to a cap domain. The ^N^RTD jelly roll was found to be structurally related to the constitutively expressed capsid protrusions (P domains) of tombusviruses, while the cap domain is absent from those viruses. A model, based on the tombusvirus structure, of how the ^N^RTD might be presented on the surface of P/E/L viruses was proposed (Schiltz et al., 2022).

To understand if and how the presence of the RTP* affects the P/E/L capsid structure and how the ^N^RTD is presented on the capsid surface, we have purified and studied two forms of TuYV particles: the wild-type virus (TuYV-WT, GenBank accession NC_003743.1, Veidt et al. (1988)), whose viral capsids contain both CP and RTP*, and a mutant (TuYV-ΔRTP) whose capsids contain the CP only. The two forms have been observed and characterised by electron microscopy (EM) techniques: negative staining, immunogold labelling and 3D cryo-EM. In this paper we present and discuss the results of our observations.

## Materials and Methods

### Virus production and purification

Purified virions were prepared from 320 g of frozen leaf material of *N. benthamiana* infiltrated with TuYV-WT full-length infectious clone (Leiser et al., 1992) using agroinfiltration (English et al., 1997) and from 335 g of *Montia perfoliata* similarly infiltrated with TuYV-ΔRTP infectious clone (referred to as BW6.4 in Brault et al (1995)). Infiltrated leaves were collected and frozen at –80 °C 6 to 8 days post-inoculation. The purification was performed according to van den Heuvel et al. (1991) and completed by a sucrose density gradient centrifugation followed by ultracentrifugation at 45 000 rpm during 2 h 30 min at 17 °C to obtain a highly concentrated suspension of virions. The final pellet was suspended in citrate buffer 0.1 M pH 6.0 and stored at –80 °C. The final concentration reached 1.32 µg/µL and 3.5 µg/µL for TuYV-WT and TuYV-ΔRTP respectively.

### Immunogold labelling of viruses

Formvar/carbon coated 300 mesh nickel grids were allowed to float for 30 min on drops of purified TuYV-WT or TuYV-ΔRTP at about 100 ng/µL in citrate buffer 0.1 M (pH 6.0). Grids were then incubated in blocking buffer (PBS + 2% BSA) for 30 min and transferred to drops containing purified immunoglobulins at 0.8 mg/mL directed against the TuYV RTD, dilution 1:40 in PBS (Reutenauer et al., 1993). Incubation was conducted overnight at 4 °C. Grids were rinsed in blocking buffer (30 min) and incubated in goat anti-rabbit IgG conjugated with 10 nm gold beads for 3 h 30 min, 1:20 in PBS 0.1% BSA (BBInternational, Cardiff, UK). Grids were rinsed with PBS then with water and finally stained with 3% uranyl acetate. Grids were observed with a Philips EM 208 transmission electron microscope operating at 80 kV.

### SDS-PAGE of purified virus

Purified TuYV-WT from *M. perfoliata* (2 µg denatured in Laemmli buffer) was loaded on a 10% SDS-PAGE which was further stained with colloidal Coomassie blue. After staining, the bands were quantified using ImageJ (see Web references). The net electrostatic charge of each structural protein was calculated using Prot Pi (see Web references).

### Negative stain electron microscopy

Negative stain EM grids were prepared by applying 3 μL of virus solution previously diluted at 0.07 mg/ml to glow-discharged collodion-carbon-coated copper grids for 2 min. The excess of liquid was blotted with Whatman N4 filter paper and the grids were washed twice in 1% w/v uranyl acetate for 1 min. The excess liquid was removed, and grids were air-dried. The grids were observed using a JEOL 2200 FS electron microscope operating at 200 kV in zero-energy-loss mode with a slit width of 20 eV and equipped with a 4k × 4k slow-scan CDD camera (Gatan inc.)

### Cryo-EM

Preliminary images were recorded using a JEOL 2200 FS electron microscope operating at 200 kV in zero-energy-loss mode with a slit width of 20 eV and equipped with a 4k × 4k slow-scan CDD camera (Gatan inc.). We initially prepared frozen-hydrated viruses deposited onto Quantifoil R2/2 or R1.2/1.3 grids (Quantifoil Micro tools GmbH, Germany) or glow discharged Lacey grid (Ted Pella Inc., USA), blotted for 3 s and then flash frozen in liquid ethane using the semi-automated plunge freezing device Vitrobot IV at 100% relative humidity. Observations revealed that even at high nominal concentrations, up to 3.5 mg/mL, the final concentration of viruses within the grid holes was not enough for an automatic image acquisition. Indeed, most of the virus particles interacted with the carbon film and TuYV-ΔRTP also formed nanocrystals. To overcome this problem, we used Quantifoil R2/2 grids covered with a 10-15 nm carbon film, which led to virus concentrations suitable for a single-particle cryo-EM analysis. No deformation of viral particles was observed, indicating that they were apparently not impaired by the interaction with the carbon film.

For high-resolution data collection, 3 μL of purified viral particles at ∼1 mg/mL were applied during 2 min to glow-discharged Quantifoil R 2/2 grids covered with a thin carbon film (Quantifoil Micro tools GmbH, Germany), blotted for 3 s and then flash frozen in liquid ethane using the semi-automated plunge freezing device Vitrobot IV at 100% relative humidity. A Polara transmission electron microscope (Thermo Fisher Scientific, Eindhoven, Netherlands) (IBS, Grenoble, France) was used operating at 300 kV with defocus ranging from –0.8 to –2.5 µm. The pixel size estimated at the specimen level was 0.97 Å. A total of 151 and 179 movies were recorded for TuYV-WT and TuYV-ΔRTP respectively, using the Gatan Latitude single particle acquisition software (Gatan inc.) on a K2 camera in counting mode. The total exposure time was 5 s corresponding to a dose of 50 e^−^/Å^2^. Forty individual frames were collected with an electron dose of 1.0 e^−^/Å^2^ per frame.

### Image processing and 3D reconstructions

Image processing was carried out using RELION 3.1 (Scheres, 2012; Zivanov et al., 2018) for icosahedral reconstructions, RELION 4.0 (Kimanius et al., 2021) for focussed asymmetric classifications, Gctf (Zhang, 2016) for contrast transfer function (CTF) calculations, Gautomatch (developed by Kai Zhang at the MRC Laboratory of Molecular Biology, UK) for automatic particle picking and RELION 3.1 for local resolution estimation. EMAN2 (Tang et al., 2007) was used to generate radial density plots.

The frames of each movie were computationally corrected for drift and beam-induced movement (Zivanov et al., 2018) and the CTF calculated. 51 930 TuYV-WT and 26 519 TuYV-ΔRTP particles were automatically picked, extracted from the movies and rescaled (pixel size: 2.91 Å, box size: 160 pixels) and submitted to reference-free 2D classification with regularisation parameter *τ*=2 and mask diameter=377 Å. 40 780 TuYV-WT and 18 433 TuYV-ΔRTP particles were selected from the best 2D classes and used to compute *de novo* 3D models at 15 Å resolution. Icosahedral symmetry was imposed during the model computation. Further particles were discarded based on the results of 3D classifications (pixel size: 1.22526 Å, box size: 380 pixels, *τ*: 4, mask diameter: 375 Å, number of classes: 4) and 3D alignment refinements with icosahedral symmetry constraints. The sets of particles producing the best reconstructions were used for per-particle defocus refinement and Bayesian polishing. Ultimately, 17 316 TuYV-WT and 3 003 TuYV-ΔRTP selected particles yielded icosahedral reconstructions at 4.1 Å and 3.5 Å resolution respectively. The resolution was estimated based on a 0.143 gold-standard FSC threshold on masked maps (Chen et al., 2013; Scheres and Chen, 2012). The refined maps were post-processed through model masking and *B-*factor sharpening, with *B* = −137 Å^2^ and –60.5 Å^2^ for TuYV-WT and TuYV-ΔRTP respectively.

To reconstruct the ^N^RTD from the TuYV-WT images, several asymmetric, focussed 3D classifications were run. Regions surrounding an *I*_2_, a *Q*_2_ and a *Q*_3_ axis of the CP-shell were independently targeted (Fig. S1; see Supplementary text for the definition of the symmetry axes symbols). These regions were chosen based on the observation that, *in vitro*, the ^N^RTD forms symmetric dimers (Schiltz et al., 2022), and that the homologous dimeric domains in tomato bushy stunt virus (TBSV, PDB 2tbv) lie on the capsid surface in strict contact with it and aligned with the capsid *I*_2_ or *Q*_2_ axes. Also, in almost all other known *T*=3 virus structures possessing protruding domains, those domains are in a dimeric arrangement around the capsid symmetry axes and in contact with the capsid surface. This has been observed in some nodavirids (PDB 6ab6) and calicivirids (PDB 3m8l, 2gh8, 7bjp; 7dod; 9cvg; 6otf; 3j1p), except for Orsay virus (PDB 4nwv), where the protruding domains are better described as trimers. We also targeted regions further away from the capsid surface to account for the length of the CP-^N^RTD linker (Fig. S1). For each targeted axis, three spherical masks were used possessing a diameter of 80 Å, 120 Å and 180 Å (Fig. S1), corresponding respectively to 1, 1.5 and 2 times the size of the ^N^RTD (estimated based on PDB 7uln). These masks were positioned either assuming the ^N^RTD in contact with the capsid surface or at a distance, estimated by assuming extended CP-^N^RTD linkers (21 residues) in polyproline helix conformation parallel to the CP symmetry axis. An elongated mask, consisting of approximately one half of an ellipsoid (half-axis lengths 50 Å, 50 Å and 180 Å), was also used (Fig. S1). Finally, an approximately hemispherical mask (radius 180 Å, Fig. S1) covering a large region around the targeted axis was used to account for the possibility of ^N^RTD locations far from the symmetry axis (bending of the linkers). A 19.6 Å (16 pixels) soft border was added to all masks. The set of 17 316 aligned images of the TuYV-WT particles was symmetry-expanded using either all 60 icosahedral-symmetry operators (for 3D classifications focussed around the *Q*_2_ and *Q*_3_ axes) or a 30-operator subset (for 3D classifications focussed around the *I*_2_ axis). Projections of the TuYV-WT reconstructed capsid were subtracted from the images of the expanded set, except for a small region of the protein shell corresponding approximately to the volume of four CP monomers surrounding the targeted *I*_2_ or *Q*_2_ axis, or three CP monomers surrounding the targeted *Q*_3_ axis. The subtracted images were recentred at the centre of the spherical masks or at the base of the hemispherical or hemi-ellipsoidal masks and re-boxed (box size: 216 pixels for spherical masks, 312 pixels otherwise; pixel size: 1.22526 Å). With each mask, extensive 3D focussed (fixed particles), asymmetric classifications were carried out (parameters: *τ*=4,10,20,40,60; number of classes=4; mask diameter=300-360 Å). No preestablished atomic model of the ^N^RTD was used as a reference for classification. All attempts to identify the ^N^RTD in the 3D classes were unsuccessful.

### Atomic model building, refinement and analysis

The TuYV-ΔRTP capsid structure was built and refined starting from an AlphaFold-3 (Abramson et al., 2024) prediction of the CP domain (Fig. S2). To save computation time and memory and, at the same time, correctly avoid steric clashes between symmetry-related neighbouring chains, only a part of the whole *T*=3 icosahedral capsid density was considered, which consisted of eleven subunits: the three chains of an asymmetric unit plus the eight closest neighbouring chains. Eleven copies of the AlphaFold-3 CP model (from which the N-terminal Arg-rich stretch was excluded) were fitted and iteratively refined against the experimental density map – following the general guidelines described by Afonine et al. (2018a) – by alternated steps of automatic refinement under stereochemical restraints with PHENIX.real_space_refine (Afonine et al., 2018b), visual model inspection and correction with *Coot* and model geometry validation with MolProbity (Williams et al., 2018). Non-crystallographic symmetry (NCS) restraints were applied during the first refinement run and subsequently released. Standard stereochemical restraints and interactively-checked secondary-structure restraints were applied during all refinement steps. Atomic isotropic displacement parameters (one parameter per residue) were applied and refined in the last refinement run. Finally, the refined model was pruned back to one asymmetric unit (three chains) before deposition with the PDB. The TuYV-WT atomic structure was similarly refined using the TuYV-ΔRTP structure as a starting point and no NCS restraint. Subunit-subunit interfaces were analysed using PISA (Krissinel and Henrick, 2007). The directions and positions of the local symmetry axes in the final structures were calculated by least-squares optimisation, under matrix-orthogonality constraints, of the group of matrices associated to an exact *n*-fold axis with respect to the transformations obtained by pair-wise alignment of the relevant subunits. The geometrical parameters describing the ideal virus architecture (Fig. S3) were calculated based on Fig. S4, basic trigonometry and the aid of formulas for regular polyhedra (icosahedron, dodecahedron and pentagonal pyramid) available from https://mathworld.wolfram.com (see Web references). Results of cumbersome trigonometric expressions were checked with the aid of Wolfram|Alpha (see Web references). The expected volumes of the unmodelled portions of the structure at the capsid interior (CP N-ter stretches and genomic RNA) were estimated using NucProt (Voss and Gerstein, 2005).

### Artwork

Figures were prepared using: UCSF Chimera (Pettersen et al., 2004), developed by the Resource for Biocomputing, Visualization, and Informatics at the University of California, San Francisco, with support from NIH P41-GM103311; UCSF ChimeraX (Meng et al., 2023), developed by the Resource for Biocomputing, Visualization, and Informatics at the University of California, San Francisco, with support from National Institutes of Health R01-GM129325 and the Office of Cyber Infrastructure and Computational Biology, National Institute of Allergy and Infectious Diseases; PyMOL open source, version 2.5.0 (Schrödinger, LLC, 2024).

## Results and Discussion

### Characterisation of purified viral particles by negative staining, immunogold labelling and SDS-PAGE

TuYV virions were purified by sucrose density centrifugation from agroinfiltrated leaves of plants inoculated with the wild-type (TuYV-WT) or a mutant clone (TuYV-ΔRTP) devoid of RTP* (Fig. 1). The mutant (called BWYV-6.4 in Brault et al. (1995)) is affected in plant systemic infection and is not transmissible by aphids (Brault et al., 1995). TuYV-ΔRTP virions are solely composed of CP subunits whereas the wild-type virions contain both CP and RTP* (Brault et al., 1995) (Fig. 1).

Observation of negatively stained TuYV-WT virions by transmission EM revealed circular particles with a diameter of approximately 30 nm (Fig. 2a). In the TuYV-ΔRTP samples two particle shapes are reproducibly observed: circular, sized as the wild-type virions (Fig. 2b), and slightly elongated (Fig. 2c). This latter shape has never been observed in preparations of purified TuYV-WT. TuYV-ΔRTP particles were rather smooth (Fig. 2b), while TuYV-WT virions displayed protruding densities at their surface (4 to 6 protrusions on each particle), indicated by black arrows in Fig. 2a, likely the ^N^RTD from the RTP*.

**Fig. 2.**
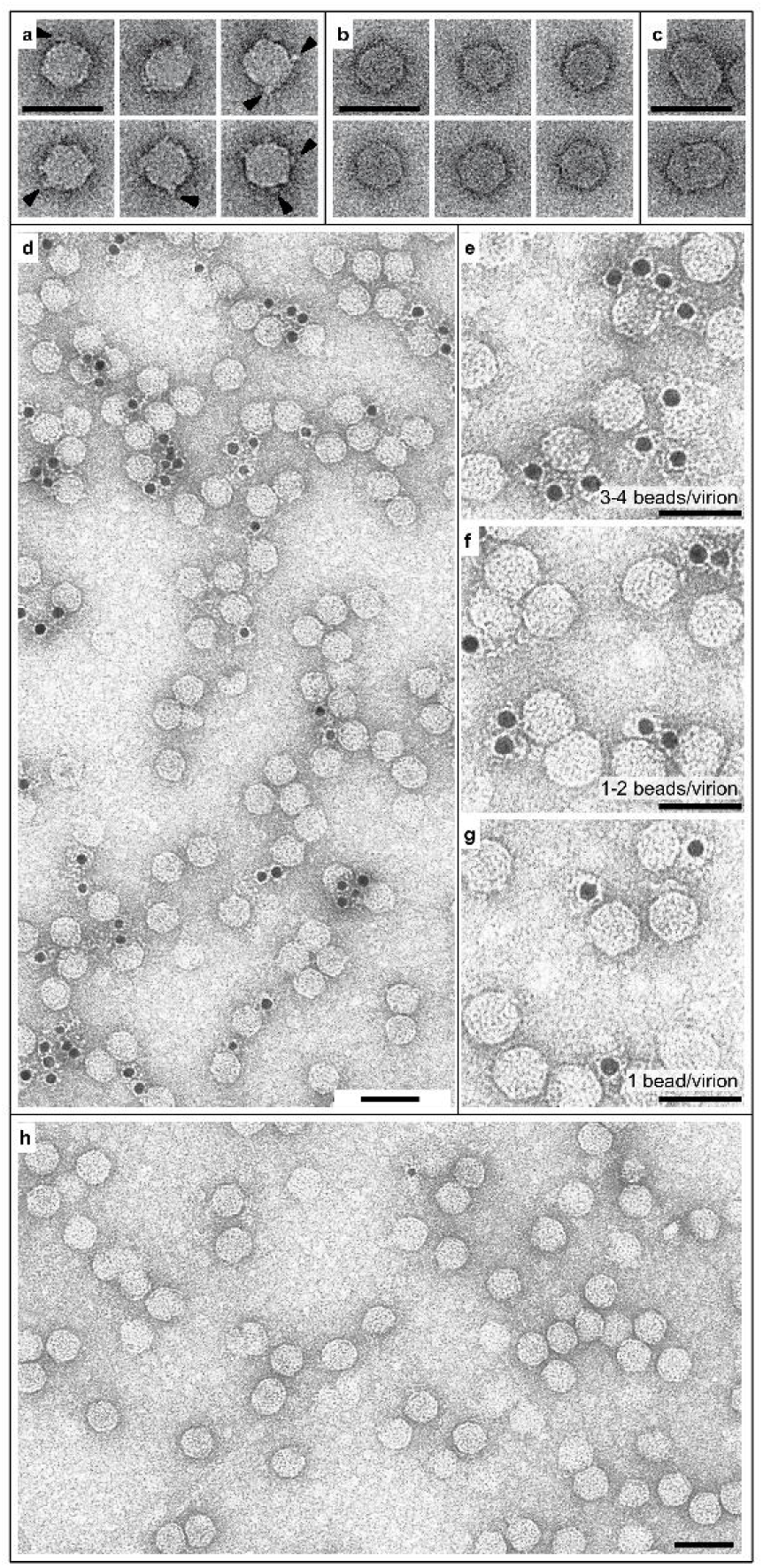
EM images of negatively stained and immunogold-labelled TuYV samples. (**a**) Negatively stained wild-type virions. Protrusions of the capsid (probable presence of the RTP*) are indicated by arrows. (**b**) Negatively stained mutated virions. No clear protrusion of the capsid is observed. (**c**) Elongated particles observed in samples of the mutant. (**d**-**g**) Immunogold-labelled wild-type virions. Gold particles (darker spots) indicate the presence of the RTP*. (**h**) Mutated virions after immunogold labelling. No gold particle is observed, indicative of the absence of the RTP*. Scale bars are 50 nm.

Immunogold labelling experiments using antibodies directed against the RTD (Reutenauer et al., 1993) confirm exposition of the ^N^RTD at the surface of the TuYV-WT virions (Fig. 2d-g) as previously shown on virions partially purified from TuYV-infected plant protoplasts (Brault et al., 1995). No gold labelling was observed on TuYV-ΔRTP virions (Figure 2f). We estimated that the number of gold beads per viral particles in the EM images (Table S1) varies from 0 to 4. This is consistent with the presence of the protrusions observed in negatively stained samples.

The number of incorporated RTP*s per TuYV-WT particle, estimated by electrophoretic analysis, was higher than the average number of beads observed by immunogold labelling and capsid protrusions observed by negative staining. Based on colloidal-Coomassie-blue SDS-PAGE band intensity (Fig. S5), the CP/ RTP* ratio was estimated to be 13.5, corresponding to 12.4 incorporated RPT*s per virion. Note, however, that estimations from stained gels are approximate (Tal et al., 1985).

Schiltz et al. (2022) showed that, *in vitro*, the ^N^RTD forms dimers. The discrepancy we observed between the number of incorporated RTP*s estimated by negative staining, immunogold labelling and electrophoretic analysis on denatured proteins is in line with the hypothesis that the observed capsid protrusions contain ^N^RTD dimers. The varying number of beads per virion observed after immunogold labelling suggests a heterogeneous population of viruses incorporating a different number of RTP* subunits. One cannot exclude, though, the lack of accessibility of some ^N^RTDs, or the presence of different RTP* conformations, some of which would not be recognised by the antibodies.

We found that TuYV-ΔRTP can form large 2D-arrays of particles in negatively stained samples (Fig. 3a) and also small 3D crystals in frozen hydrated samples at a concentration of 3.5 mg/mL (Fig. 3b) as already seen in aphids fed on purified TuYV-ΔRTP (Reinbold et al., 2001). The regularity of these assemblies seems to be interrupted by the incorporation of elongated particles, indicated by arrows in Fig. 3a. This extensive regular organisation has never been observed for the wild-type virions neither by negative stain nor by cryo-EM. Only some assemblies of 2 or 4 virions were observed in negatively stained samples (Fig. 3c). This different behaviour suggests that the presence of RTP* in TuYV-WT strongly impairs the interplay between viral particles.

**Fig. 3.**
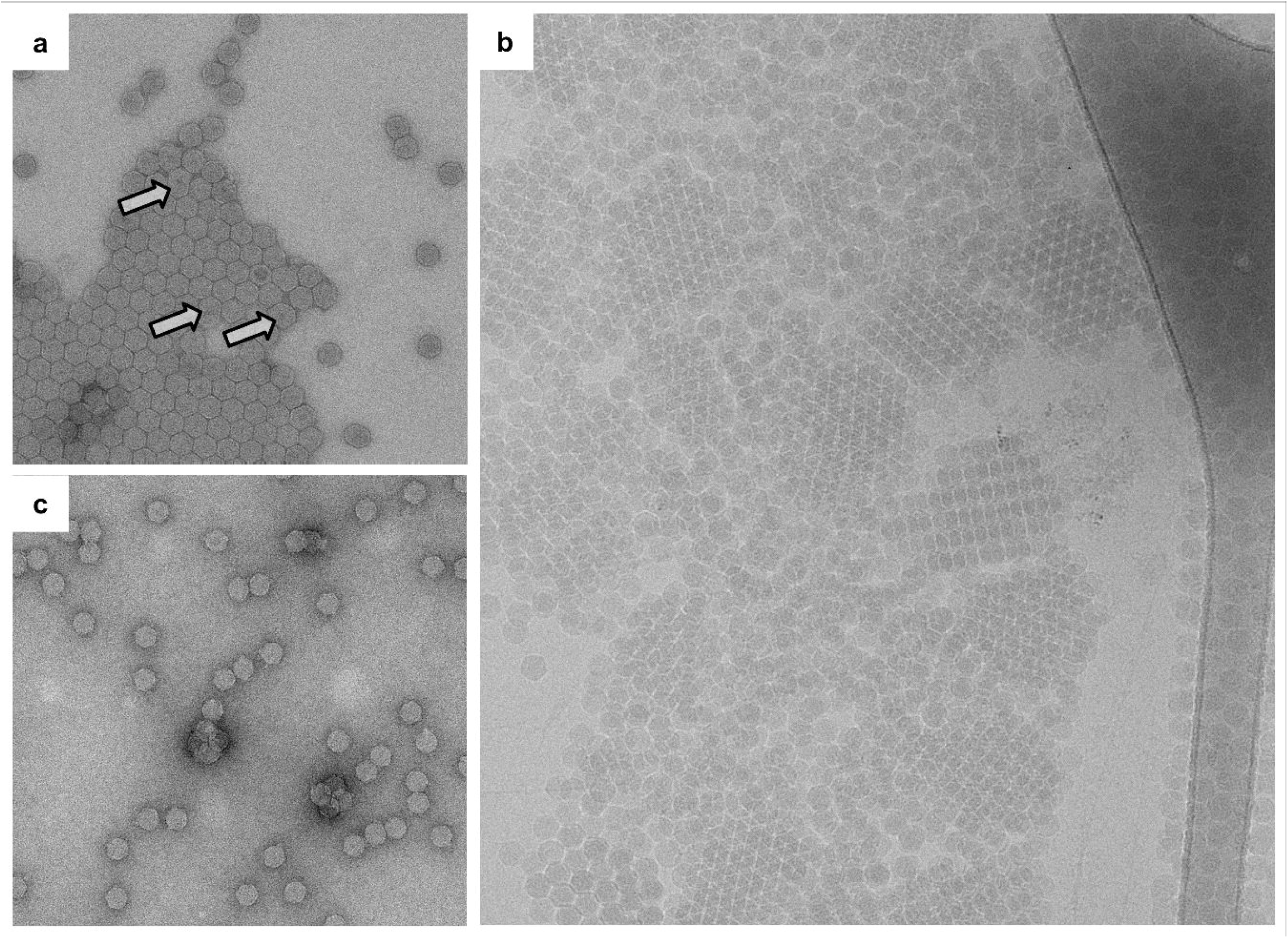
TuYV-ΔRTP assembles into nanocrystals, while TuYV-WT does not. (**a**) Two-dimensional array of negatively stained TuYV-ΔRTP virions. Arrows indicate elongated or distorted particles. (**b**) Cryo-EM image of TuYV-ΔRTP particles arranged in small, thin 3D crystals. (**c**) Negatively stained TuYV-WT virions forming small assemblies of 2 to 4 particles.

### Cryo-EM 3D reconstruction of TuYV-WT and TuYV-ΔRTP

Frozen-hydrated samples of TuYV-WT and TuYV-ΔRTP were prepared on Quantifoil R2/2 holey-carbon grids covered with a carbon film to avoid preferential aggregation at the hole borders and formation of nanocrystals (Fig. 3b). Transmission EM images of the samples were collected using a Polara microscope equipped with a K2 camera. In most of the images, uniform distributions of mostly undistorted, RNA-filled spherical virions were observed (Fig. 4a). The 3D scattering densities of the TuYV-WT and TuYV-ΔRTP virions were reconstructed from selected sets of, respectively, 17 316 and 3 003 homogeneous, RNA-filled particles (Fig. 4b-c). Statistics for the cryo-EM single-particle analysis are summarised in Table S2. The overall resolution of the reconstructed maps, refined by imposing icosahedral symmetry constraints, was respectively estimated to be 4.1 Å and 3.5 Å for TuYV-WT and TuYV-ΔRTP (0.143 gold-standard FSC criterion). Both the protein shell, which was well resolved in the sharpened maps (Fig. 4b) and the RNA shell obscured by icosahedral averaging, are visible in the unsharpened, unmasked maps (Fig. 4c). Local resolution in the protein shell varies in the range 3.95–4.5 Å and 3.3–4.5 Å for TuYV-WT and TuYV-ΔRTP respectively (Fig. S6). Atomic models of the capsids, comprising more than two thirds of the length of the CP domains (residues 62/64 to 202), have been built and refined against the reconstructed densities (Fig. 4d-h, Fig. S7, Table S3). No interpretable density could be attributed to the long arginine-rich N-terminal stretches (Fig. 4h), which are predicted to be disordered and likely buried in the nucleic acid at the interior of the virions. Expectedly, due to symmetry averaging during the reconstruction procedure, no extra density indicative of the presence of the ^N^RTD shows up in the TuYV-WT reconstruction (asymmetric reconstructions are discussed later). The atomic models of the CP assemblies do not reveal any significant difference between the TuYV-WT and TuYV-ΔRTP protein shells (Fig. 4e). Direct comparison of the trimers in the icosahedral asymmetric unit, without rigid-body adjustments of their positions, shows a low root mean square deviation (RMSD) between main-chain atoms (0.39 Å). Rigid body fit of individual subunit pairs results in slight improvements of the RMSD (0.32-0.34 Å), but with negligible movements of the subunits (0.2-0.6° rotations).

**Fig. 4.**
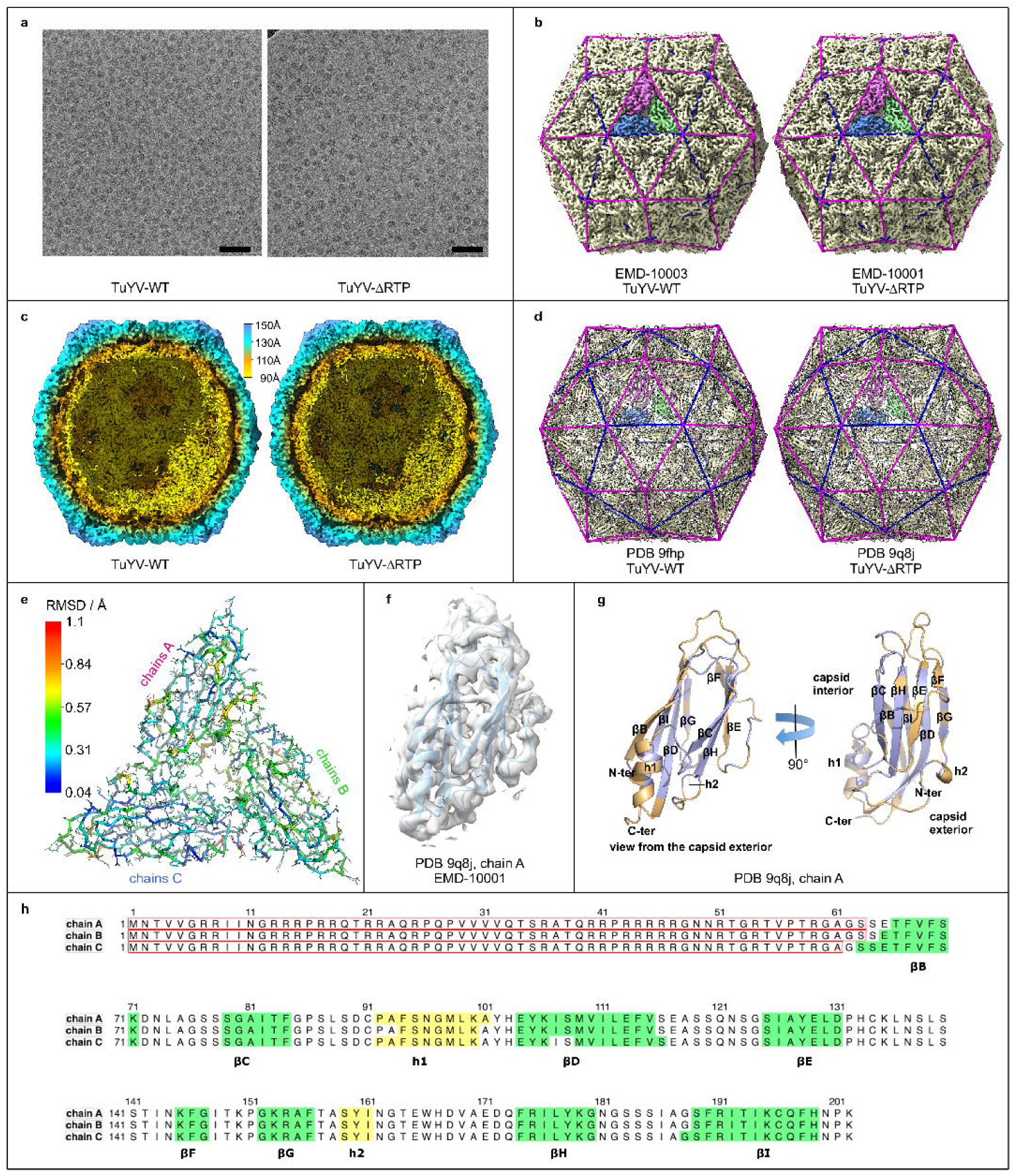
Cryo-EM reconstruction of TuYV-WT and TuYV-ΔRTP. (**a**) Motion-corrected EM images of frozen-hydrated virions deposited on Quantifoil R2/2 grids covered with a carbon film. Scale bar is 100 nm. Images are low-pass filtered at 15 Å for illustration purposes. (**b**) 3D scattering-density (sharpened maps) of the TuYV virions reconstructed by imposing icosahedral symmetry. Three protein subunits, composing one asymmetric unit of the assembly, are highlighted in colours (blue, magenta, green). The underlying geometry of the assembly (equilateral elevated dodecahedron) is represented by coloured (blue and magenta) edges. The EMDB accession codes are reported. (**c**) Sectional view of the unsharpened, unmasked maps, illustrating the outer (protein) and inner (RNA) layers. The density is coloured according to the distance from the centre of the particle. (**d**) Atomic models of the capsid protein layers. (**e**) Comparison of the TuYV-WT and TuYV-ΔRTP atomic models. The icosahedral asymmetric units (trimers) are shown superposed on to each other and coloured by per-residue RMSD. (**f**) Ribbon representation of the atomic model of one of the CP subunits fitted in the cryo-EM map. (**g**) Two views of the atomic model of one of the CP subunits. Secondary structure elements are labelled. Residues involved in inter-subunit contacts are coloured orange. (**h**) Sequence of the three quasi-equivalent CP domains. Unmodelled regions (no interpretable density) are within red frames. Secondary structure elements are highlighted and labelled as in (g).

### The capsid architeture

If not otherwise stated, the terms “chain” and “subunit” in this subsection will refer to the visible (modelled) part of the CP domains in the TuYV capsid structures (unframed sequences in Fig. 4h).

The TuYV capsid is composed by 180 copies of the CP domain arranged according to global icosahedral rotational symmetry (symmetry axes denoted *I_n_*, where *n*=2,3,5 is the symmetry axis order) with triangulation number *T*=3, and a set of local symmetry axes (denoted *Q_n_*, *n*=2,3,5,6) (Fig. 4b,d, Fig. S3, Fig. S4). One icosahedral asymmetric unit comprises three quasi-equivalent subunits related by a *Q*_3_ axis. Each subunit folds into a classical jelly roll (Fig. 4g), commonly found in viruses, composed of two packed four-stranded anti-parallel β-sheets. In TuYV, one of the β-sheets faces the interior of the capsid while the other faces the exterior. Two intervening helices, most of the loops, the N- and C-ter β-strands and a few residues from other strands of the jelly roll provide interactions between subunits (Fig. 4g). Interacting residues at the subunit interfaces are mainly of a polar or charged nature (Fig. S8, Table S4, Table S5).

The structure of the TuYV capsid shows no significant differences with the previously published structures of polerovirus VLPs (Byrne et al., 2019) (RMSD ∼1.0 Å in comparison with PLRV and BYDV VLPs). More generally, the *T*=3 subunit arrangement of TuYV follows the schematic initially described by Rossmann et al. (1983a, 1983b) for southern bean mosaic virus (SBMV, species *Sobemovirus SBMV*), in which the quasi-equivalence of contacts (Caspar and Klug, 1962) is assured essentially by different relative orientations of the subunits at the *Q*_2_ and *I*_2_ interfaces (dimeric interfaces) where the h1-helices touch each other. The change in subunit orientation maintains the two helices in close contact but disrupts interactions and separate the underlying N-ter and C-ter β-sheets in adjacent chains (Fig. S8). Differently to what happens in SBMV, the cryo-EM maps of TuYV indicate that the increased space available under the h1-helices at the *I*_2_ interfaces is not occupied by an additionally ordered stretch of polypeptide chain.

The geometry of the overall assembly can be described, within the framework of Caspar and Klug’s theory on the principles governing regular virus structures (Caspar and Klug, 1962), in terms of an equilateral elevated dodecahedron (EED), outlined in Fig. 4b,d and described in Supplementary text, Fig. S3 and Fig. S4 (but see also Brinkmann et al. (2017), Janner (2006a, 2006b), Keef et al. (2013), Konevtsova et al. (2024), Lorman & Rochal (2008), Mannige & Brooks (2008, 2009, 2010), Rochal et al. (2017), Twarock (2006) and Twarock & Luque (2019) for different or more general approaches to the description of capsid architectures). Each triangular face or the polyhedron represents, with respect to the icosahedral symmetry, one asymmetric unit. The dihedral angles between triangular faces sharing a common edge are convex towards either the capsid interior or the capsid exterior, so that two dihedral angle values, named respectively *endo* and *exo*, and two classes of edges (named accordingly) can be defined (Fig. S3c). The EED geometrical model can be applied, to different degrees of approximation, to many but not all viruses with *T*=3 icosahedral symmetry. TuYV shows small deviations from this ideal geometry (Fig. S3g,i). We have analysed all virus structures classified as *T*=3 in the VIPERdb (Montiel-Garcia et al., 2021) and found that, among those that can be reconducted to the EED geometry, a very limited number strictly adhere to that model, for example panicum mosaic virus (species *Panicovirus panici*), while most of the others, like tomato bushy stunt virus (TBSV, species *Tombusvirus lycopersici*) and SBMV, deviate to some extent (imperfect *Q*_3_ rotations within the asymmetric unit). Other geometrical descriptions have been used in the past for some of these viruses. TBSV and SBMV have been described as rhombic triacontahedra (Johnson and Olson, 2021; Winkler et al., 1977), polyhedra composed by non-equilateral triangles forming, in pairs, flat rhombic faces (*exo* angle = 180°). The polerovirus PLRV has also been described as possessing flat *exo* angles (Adams et al., 2022), but by comparing it with TuYV we propose that PLRV also has non-flat *exo* interfaces. Other *T*=3 icosahedral viruses, like brome mosaic virus (species *Bromovirus BMV*) cannot be described by the EED model. Interestingly, none of the known *T*=3 structures adheres to the geometry of a regular icosahedron (Fig. S4).

### The RNA structure

The TuYV genomic RNA shows up in the unmasked, unsharpened maps as an amorphous layer of averaged density just underneath the CP shell (Fig. 5). The approximate volume of the residual density layer appears to be adequate to accommodate the whole genomic RNA plus the unmodelled N-ter stretches of the CP subunits (Fig. S9). Highest values of the residual density are found around the *I*_5_ and *Q*_3_ axes and underneath the BC and AC interfaces (Fig. 5b-e). CP residues in proximity of the residual density are highlighted in Fig. 5f-h.

**Fig. 5.**
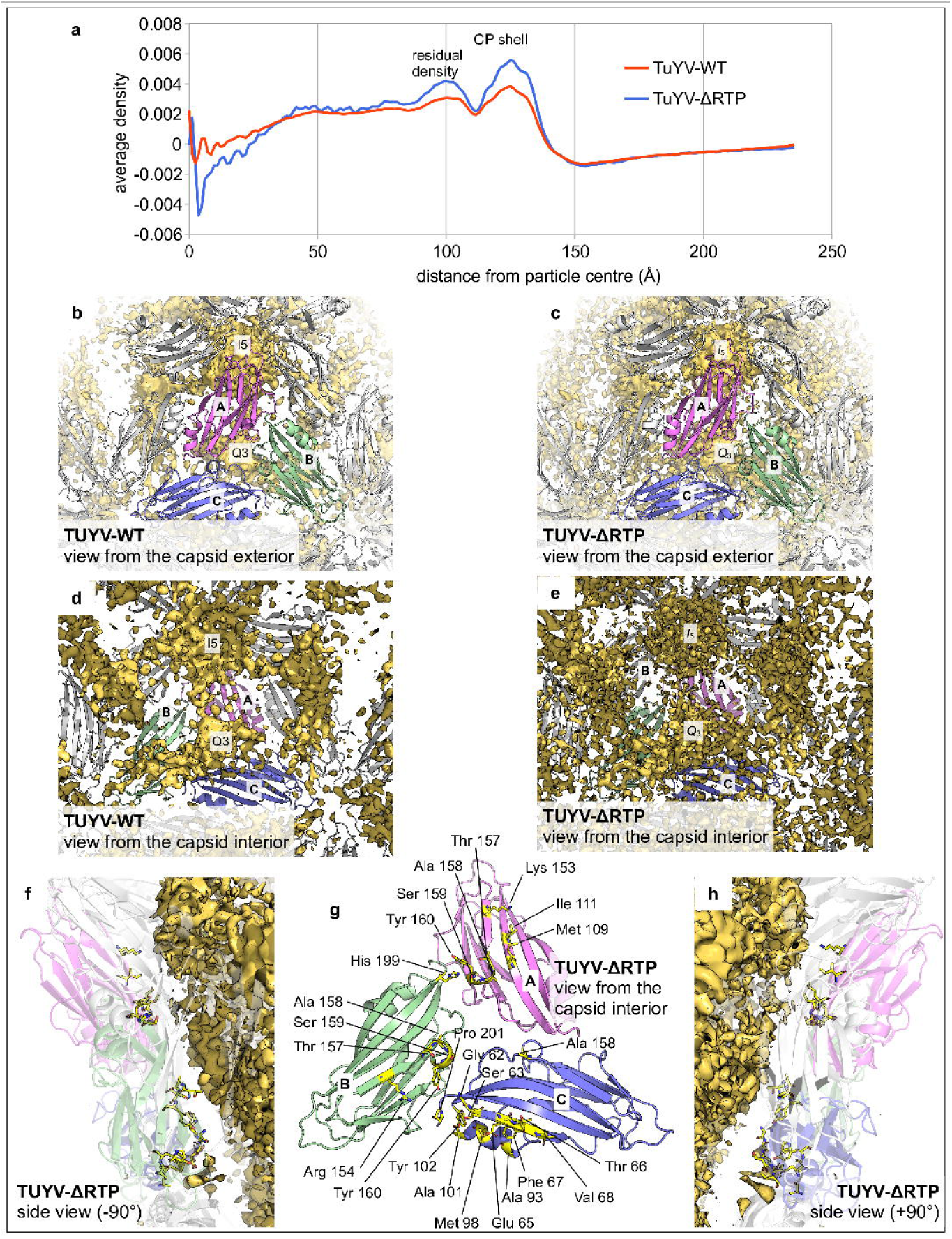
The residual amorphous density. (**a**) Radially averaged density in the unsharpened, unmasked cryo-EM maps. (**b-e**) The residual density (yellow surfaces) after masking out the regions occupied by the CP subunits (shown as cartoon models). A spherical mask of radius 3 Å has been applied to each modelled CP atom. The densities have been contoured at a level corresponding to the peak of the CP-shell in the radially averaged density (the second peak in (**a**)). (**f**-**h**) Residues of the TuYV-ΔRTP atomic model found in proximity (distance ≤ 5 Å) of the contoured residual density. Similar results (not shown) have been found for TuYV-WT.

### The unreconstructed ^N^RTD is likely to float at the capsid exterior

Inspection of some selected cryo-EM images confirmed the presence of the ^N^RTD (four to six protrusions of the capsid) in the purified TuYV-WT samples (Fig. 6), but its scattering density was not visible in the cryo-EM map of the capsid due to the symmetry averaging. No differences were observed between the radially averaged densities of the TuYV-WT and TuYV-ΔRTP particles that could be related to the ^N^RTD (Fig. 5a), indicating that the data at our disposal do not allow the 3D reconstruction of the ^N^RTD domain. Indeed, no convincing result could be obtained from focussed, asymmetric 3D classifications procedures, despite our extensive attempts (see Materials and Methods).

**Fig. 6.**
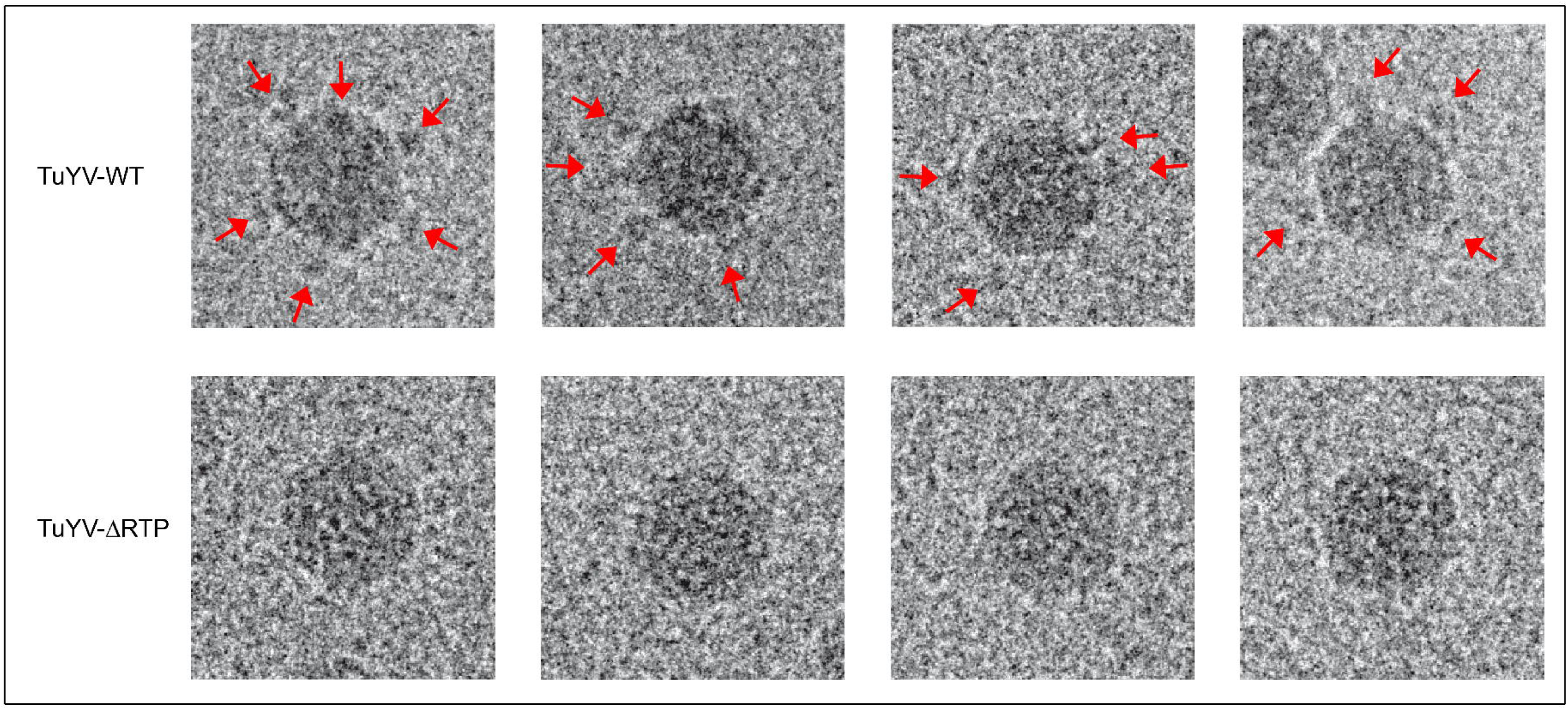
Observation of capsid protrusions derived from ^N^RTD on TuYV-WT virions by cryo-EM. Top: cryo-EM images of purified TuYV-WT virions in which some ^N^RTDs, indicated by arrows, are visible in the proximity of the capsids. Bottom: for comparison, some images of the TuYV-ΔRTP in similar orientations.

The ^N^RTD dimers described by Schiltz et al. (2022) are structurally related to the dimeric P domains that protrude along the *I*_2_ and *Q*_2_ axes of the capsids of TBSV and other tombusviruses. Based on the TuYV-ΔRTP structure described in this paper (released by the PDB in July 2022), and by comparison with TBSV, Schiltz et al. (2022) showed that, in principle, it is structurally possible for the ^N^RTD dimers to be accommodated (like in TBSV) in tight contact with the TuYV CP layer and even occupy the complete set of 90 twofold sites on the capsid surface. Failure of our extensive attempts to reconstruct the ^N^RTD around the twofold sites suggests, however, that the ^N^RTD domain does not occupy (in the conditions of our purified samples) a well-defined position on the capsid surface. Indeed, the proline-rich linker (21 residues) connecting the CP and ^N^RTD domain in TuYV (Fig. S2) is much longer than the stretch (7 residues) joining the TBSV P domain to the capsid shell, and this may allow the ^N^RTD to float around, at least in absence of a specific interacting partner, in proximity of the capsid surface.

We have seen no significant differences between the cryo-EM maps of TuYV-WT and TuYV-ΔRTP, indicating that the presence of the ^N^RTD domains did not influence the final CP fold and assembly organisation (of the selected particles). One cannot exclude, however, the possibility of ^N^RTD playing a role in RNA packaging and capsid assembly (formation of elongated particles has been observed in TuYV-ΔRTP but not in TuYV-WT samples).

## Conclusions

We presented here the structural characterisation by immunogold labelling and 3D cryo-EM of the wild-type TuYV and a mutant whose capsid contains the CP only. We showed that incorporation of the RTP* does not impair the capsid structure, and the ^N^RTD does not adopt well-defined positions at the capsid surface. The varying number of beads per virion observed after immunogold labelling suggests a heterogeneous population of viruses incorporating a different number of RTP* subunits, though one cannot exclude the lack of accessibility of some ^N^RTDs, or the presence of different RTP* conformations, some of which would not be recognised by the antibodies. This raises the question whether different forms exist of the ^N^RTD at the surface of the capsid that may have different cellular partners and fulfil different functions in the virus cycle. Answers to these questions will require further investigation.

## Deposited data references

The cryo-EM maps of TuYV-WT and TuYV-ΔRTP have been deposited with the Electron Microscopy Data Bank and released under the accession IDs EMD-10003 and EMD-10001 respectively. The model atomic coordinates have been deposited with the Protein Data Bank and released under the PDB IDs 9fhp and 9q8j respectively.

## Credits

**Joséphine Lai-Kee-Him**: Investigation, Resources. **Stefano Trapani**: Conceptualisation, Data Curation, Formal analysis, Investigation, Software, Validation, Visualisation, Writing - Original Draft, Writing - Review & Editing. **Sylvaine Boissinot**: Investigation, Resources, Writing - Review & Editing. **Catherine Reinbold**: Investigation, Resources, Visualisation, Writing - Review & Editing. **Chloé Fallet**: Investigation. **Aurélie Ancelin**: Investigation. **François Lecorre**: Investigation. **François Hoh**: Investigation. **Véronique Ziegler-Graff**: Conceptualisation, Writing - Review & Editing. **Véronique Brault**: Conceptualisation, Investigation, Project administration, Writing - Original Draft, Writing - Review & Editing. **Patrick Bron**: Conceptualisation, Funding acquisition, Investigation, Project administration, Supervision, Writing - Original Draft, Writing - Review & Editing.

## Supplementary material

Supplementary text. The EED architecture of the TuYV capsid.

Table S1. Estimation of the number of immunogold-labelled TuYV particles.

Table S2. Cryo-EM data collection and single-particle processing statistics.

Table S3. Atomic model refinement statistics.

Table S4. Geometric relationships between interacting subunits.

Table S5. Residues at the interfaces between subunits.

Fig. S1. The masks that have been used for asymmetric reconstructions of the ^N^RTD.

Fig. S2. AlphaFold-3 model of the TuYV RTP* monomer.

Fig. S3. Architecture of the TuYV capsid.

Fig. S4. Folding of a hexagonal lattice into either an icosahedron or an elevated dodecahedron.

Fig. S5. SDS-PAGE analysis and colloidal Coomassie blue staining of 2 µg of TuYV-WT viral particles purified from *M. perfoliata* and denaturated in Laemmli buffer.

Fig. S6. Local resolution of the cryo-EM reconstructions (sharpened maps).

Fig. S7. Some details of the TuYV-ΔRTP cryo-EM map and the corresponding refined atomic model.

Fig. S8. The eight non-equivalent subunit interfaces.

Fig. S9. Estimation of the residual density volumes at the capsid interior.

