## Supplemental Material for "Structure of the turnip yellows virus particles"

**SUPPLEMENTARY DATA**

**Supplementary text.**
**The EED architecture of the TuYV capsid.**

If not otherwise stated, the terms "chain" and "subunit" will hereafter refer to the visible (modelled) part of the CP domains in the TuYV capsid structures (unframed sequences in Fig. 4h). Though we report calculated values (e.g. RMSD, rotation angles) for the TuYV-ΔRTP structure only, all the considerations that follow are valid for both TuYV-WT and TuYV-ΔRTP.

The TuYV capsid is composed by 180 copies of the CP domain arranged according to icosahedral rotational symmetry with triangulation number $T$=3 (Caspar and Klug, 1962). One icosahedral asymmetric unit comprises three quasi-equivalent subunits (chains A, B and C in Fig. S3a) related by an almost exact, local threefold axis, denoted $Q_3$ (Fig. S3a-b). Each subunit folds into a classical jelly roll (Fig. 4g), commonly found in viruses, composed of two packed four-stranded anti-parallel β-sheets. In TuYV, one of the β-sheets faces the interior of the capsid while the other faces the exterior. Two intervening helices, most of the loops, the N- and C-ter β-strands and a few residues from other strands of the jelly roll provide interactions between subunits (Fig. 4g).

The geometry of the overall assembly can be described in terms of an equilateral elevated dodecahedron (EED, Fig. S3c). This is a non-convex polyhedron composed of sixty equilateral triangular faces (Janner, 2006b), where each face represents, with respect to the icosahedral symmetry of the assembly, one asymmetric unit. The sixty faces of the EED, grouped by five around the twelve fivefold axes, can be thought of as the faces of pentagonal pyramids whose bases sit on the faces of an underlying regular dodecahedron (outlined by blue edges in Fig. S3c). The dihedral angles between triangular faces sharing a common edge are convex towards either the capsid interior or the capsid exterior, so that two dihedral angle values, named respectively *endo* and *exo*, and two classes of edges (named accordingly) can be defined (Fig. S3c). The actual TuYV geometry is quite close to that of an ideal EED. The values of the *endo* and *exo* angles (141° and 174° respectively), evaluated by comparing the directions of the TuYV $Q_3$ axes, are close to the ideal values (138.19° and 168.68° respectively) (Fig. S3f-g). The virtual hinge movement that would transform an *exo* into an *endo* interface corresponds to a rotation of ~45° in TuYV as compared to 53.13° in an ideal EED.

Besides the global icosahedral symmetry, the EED arrangement implies a set of local symmetry axes. Fig. S3d and Fig. S4 illustrate how the EED global and local symmetry can be derived by virtually folding an ideal 2D hexagonal lattice according to the *endo* and *exo* angles. We have used the symbols $I_n$ ($n$ = 2, 3, 5) and $Q_n$ ($n$ = 2, 3, 5, 6) to denote, respectively, global (icosahedral) and local (quasi) $n$-fold symmetry. The approximate positions of the $I_n$ and $Q_n$ axes are indicated in Fig. S3e. The TuYV capsid can be formally described as an assembly of pentamers and hexamers centred on the $I_5$ and $I_3$ icosahedral axes respectively. Fig. S3e shows a set of two hexamers, chains $BC_2B_3C_4B_5C_6$ and $CB_2C_3B_4C_5B_6$, and one pentamer, chains $AA_2A_3A_4A_5$. This portion of the capsid encompasses one asymmetric unit plus all its adjacent subunits. Pentamers obey strict fivefold symmetry. Hexamers obey threefold but not sixfold symmetry and are composed of two sets of interleaved subunits, blue and green subunits respectively in Fig. S3h, not related by icosahedral symmetry. Large deviations from ideal sixfold symmetry are observed around the $I_3$ axes. Superposition of adjacent chains within a hexamer requires either a quasi-sixfold ($Q_6$) rotation of 60.2° or a quasi-fivefold ($Q_5$) rotation of 67.3° around axes that are not collinear with the $I_3$ axis. The $Q_6$-$I_3$ and $Q_5$-$I_3$ axes misalignments are 23.0° and 20.7° respectively (Fig. S3i). Note that $Q_5$ rotations give rise to an arrangement of adjacent subunits (ex. subunits $CB_2$ in Fig. S3e,h,i) similar to that observed within pentamers (ex. subunits $AA_2$ in Fig. S3e), hence the preferred symbol $Q_5$ instead of $Q_6$. Also, non-equivalent triplets of subunits can be found (e.g. $ABA_2$ and $CAB_2$) which are arranged essentially in the same way and whose local symmetry axes would be compatible with those of a $T$=1 icosahedron (Fig. S3j, Fig. S8). This is a consequence of pentamers and hexamers sharing *endo* angles (Fig. S3e), a structural constraint described as "*endo* angle propagation" by Mannige & Brooks (2010, 2009).

A total of eight independent —*i.e.* not related by icosahedral symmetry— interfaces between subunits exist. We have labelled the interfaces according to the rotation $I_n$ or $Q_n$ relating the contacting subunits (Table S4). The portion of capsid illustrated in Fig. S3e contains at least one representative of the eight interfaces. Interacting residues at the subunit interfaces are mainly of a polar or charged nature (Table S5, Fig. S8). No major differences are observed among the three $Q_3$ interfaces, though these are not strictly equivalent in principle, nor between the $Q_5$ and $I_5$ interfaces. (Table S5, Fig. S8). On the other hand, significant differences are found at the $I_2$, when compared to $Q_2$, and $Q_6$, when compared to $Q_5$ or $I_5$ interfaces. These differences are related to the existence, in an EED, of different dihedral angles: the $Q_2$, $Q_5$ and $I_5$ interfaces are found on *endo* edges, while the $I_2$ and $Q_6$ interfaces are found on *exo* edges. Indeed, most residues involved in $Q_5$, $I_5$, $Q_6$, $Q_2$ and $I_2$ interfaces line the edges of the EED (Fig. S8). The virtual hinge movement transforming an *endo* interface into an *exo* interface (Fig. S3g) would occur around a line of atomic contacts between helices h1 of neighbouring subunits. This transformation maintains the two helices in close contact but disrupts interactions and separate the underlying N-ter and C-ter β-sheets in adjacent chains (Fig. S8).

| Grids | Particles with no gold bead | Particles with one gold bead | Particles with two gold beads | Particles with three or four gold beads | % of labelled particles |
|---|---|---|---|---|---|
| TuYV-WT #958 | 348 (64.4%) | 165 (30.6%) | 27 (5.0%) | 0 (0%) | 35.6% |
| TuYV-WT #968 | 356 (60.8%) | 197 (33.6%) | 28 (4.8%) | 5 (0.8%) | 39.2% |
| TuYV-WT #960 | 317 (66.2%) | 129 (26.9%) | 29 (6.1%) | 4 (0.8%) | 33.8% |
| TuYV-WT #959 | 351 (67.2%) | 145 (27.8%) | 24 (4.6%) | 2 (0.4%) | 32.8% |
| TuYV-WT #966 | 351 (61.6%) | 163 (28.6%) | 44 (7.7%) | 12 (2.1%) | 38.4% |
| TuYV-ΔRTP #970 | 393 (99.0%) | 4 (1.0%) | 0 (0%) | 0 (0%) | 1.0% |
| TuYV-ΔRTP #973 | 470 (99.4%) | 2 (0.4%) | 1 (0.2%) | 0 (0%) | 0.4% |
| TuYV-ΔRTP #972 | 394 (99.5%) | 2 (0.5%) | 0 (0%) | 0 (0%) | 0.5% |
| TuYV-ΔRTP #974 | 416 (98.8%) | 4 (1.0%) | 1 (0.2%) | 0 (0%) | 1.2% |
| TuYV-ΔRTP #975 | 465 (98.9%) | 4 (0.9%) | 1 (0.2%) | 0 (0%) | 1.1% |

**Table S1. Estimation of the number of immunogold-labelled TuYV particles.**
Estimation of the number of TuYV particles labelled with different numbers of gold beads after incubation with immunoglobulins specifically directed against the TuYV RTD. Observations made on 10 samples (grids), 5 containing the wild-type and 5 containing the mutated TuYV-ΔRTP virus.

| | **TuYV-ΔRTP EMD-10001** | **TuYV-WT EMD-10003** |
|---|---|---|
| Data acquisition | | |
| Electron microscope | FEI POLARA 300 | FEI POLARA 300 |
| Direct electron detector | GATAN K2 SUMMIT (4k x 4k) | GATAN K2 SUMMIT (4k x 4k) |
| Magnification | 20 000 | 20 000 |
| Voltage (kV) | 300 | 300 |
| Micrographs collected | 179 | 151 |
| Frames per exposure | 40 | 40 |
| Exposure time (s) | 5 | 5 |
| Per-frame electron exposure | 1.0 | 1.0 |
| Defocus range (µm) | 0.8-2.5 | 0.8-2.5 |
| Pixel size (Å) | 0.97 | 0.97 |
| Processing | - | |
| Software | RELION 3.1 | RELION 3.1 |
| Extraction pixel size (Å) | 1.22526 | 1.22526 |
| Extraction box size (pixels) | 380 | 380 |
| Symmetry imposed | chiral icosahedral | chiral icosahedral |
| Refinement type | gold standard | gold standard |
| No. of initial particle images | 26 519 | 51 930 |
| No. of final particle images | 3 001 | 17 316 |
| Map resolution (Å) | 3.5 | 4.1 |
| FSC threshold (for map-resolution estimation) | 0.143 | 0.143 |
| Map-sharpening $B$ factor (Å$^2$) | –60.5 | –137 |

**Table S2. Cryo-EM data collection and single-particle processing statistics.**

| | **TuYV-ΔRTP PDB-9q8j** | **TuYV-WT PDB-9fhp** |
|---|---|---|
| Initial model used | AlphaFold-3 prediction | PDB-9q8j |
| Model resolution (Å) | 3.47 | 4.06 |
| Map/model FSC threshold (for model resolution estimation) | 0.143 | 0.143 |
| Model composition (one asymmetric unit, chains A, B, C) | | |
| Non-hydrogen atoms | 3 208 | 3 208 |
| Protein/nucleic acid | 3 208 | 3 208 |
| Ligands | 0 | 0 |
| B factors (average \| min \| max, Å$^2$) | | |
| Protein | 68.08 \| 54.85 \| 80.23 | 45.69 \| 30.59 \| 65.08 |
| Ligand | — | — |
| R.m.s. deviations from ideality (Engh, R. A. & Huber, R. (1991). Acta Cryst. A47, 392–400) | | |
| Bonds (Å) | 0.005 | 0.004 |
| Angles (°) | 0.988 | 0.950 |
| Clashscore | 5.9 | 5 |
| Poor rotamers (%) | 1.97 | 1.69 |
| Ramachandran plot (Oldfield, T. J. (2001). Acta Cryst. D57, 82–94) | | |
| Favoured (%) | 97.1 | 97.3 |
| Allowed (%) | 2.9 | 2.7 |
| Disallowed (%) | 0 | 0 |

**Table S3. Atomic model refinement statistics.**

| Superposed chains [a] | $BC \approx B_6C_2$ | AB | CA | $C_2B \approx CB_6 \approx C_3B_2$ | $A_2A \approx AA_5$ | $B_2C \approx BC_6$ | $CC_2$ | $A_2B \approx AB_2$ |
|---|---|---|---|---|---|---|---|---|
| $C^{\alpha}$ RMSD | 0.43 Å | 0.47 Å | 0.48 Å | 0.43 Å | 0.00 Å | 0.43 Å | 0.00 Å | 0.47 Å |
| Rotation angle | 119.87° | 120.30° | 119.96° | 60.19° | 72.00° | 67.34° | 180.00° | 178.78° |
| Translation amplitude | 0.33 Å | 0.45 Å | 0.13 Å | 0.87 Å | 0.00 Å | 0.58 Å | 0.00 Å | 1.28 Å |
| Rotation label | $Q_3$ | $Q_3$ | $Q_3$ | $Q_6$ [b] | $I_5$ | $Q_5$ [b] | $I_2$ | $Q_2$ |

[a] Chain labels as in Fig. S3e. The "≈" symbol indicates equivalent interactions according to icosahedral symmetry.
[b] Note that $Q_6$ and $Q_5$ compose an $I_3$ rotation: $Q_6 \cdot Q_5 = I_3$.

**Table S4. Geometric relationships between interacting subunits.**

Results of pairwise rigid-body superposition of interacting subunits found within one asymmetric unit and its neighbourhoods. Chains are labelled as in Fig. S3e.

| Interface type § | $Q_3$ | | | $Q_6$ | $I_5$ | $Q_5$ | $I_2$ | $Q_2$ |
|---|---|---|---|---|---|---|---|---|
| Subunit pairs § | BC | AB | CA | $C_2B$ | $A_2A$ | $B_2C$ | $CC_2$ | $A_2B$ |
| Interface residues † | B: Glu 129 (s) | A: Glu 129 (s) | C: Glu 129 (s) | | $A_2$: Val 68 | $B_2$: Val 68 | | $A_2$: Ser 64 (s) |
| | B: Leu 130 (h) | A: Leu 130 (h) | C: Leu 130 (h) | | $A_2$: Phe 69 | $B_2$: Phe 69 | | $A_2$: Glu 65 |
| | B: Asp 131 | A: Asp 131 | C: Asp 131 | $C_2$: Ser 70 | $A_2$: Ser 70 (h) | $B_2$: Ser 70 | | $A_2$: Thr 66 |
| | B: Pro 132 | A: Pro 132 | C: Pro 132 | $C_2$: Asp 72 (h) | $A_2$: Asp 72 (h) | $B_2$: Asp 72 (h) | C: Phe 67 | $A_2$: Phe 67 |
| | B: His 133 (hs) | A: His 133 | C: His 133 (s) | $C_2$: Asn 73 (h) | $A_2$: Asn 73 | $B_2$: Asn 73 | | $A_2$: Val 68 |
| | B: Lys 135 (s) | A: Lys 135 | C: Lys 135 (s) | $C_2$: Val 115 | $A_2$: Val 115 | $B_2$: Val 115 | | $A_2$: Cys 91 |
| | B: Leu 136 | A: Leu 136 | C: Leu 136 | | $A_2$: Ser 116 (h) | $B_2$: Ser 116 (h) | C: Pro 92 | $A_2$: Pro 92 |
| | B: Ser 140 | | C: Ser 140 | $C_2$: Glu 117 (hs) | $A_2$: Glu 117 (hs) | $B_2$: Glu 117 (hs) | C: Ala 93 | $A_2$: Ala 93 |
| | B: Ser 141 (h) | A: Ser 141 (h) | C: Ser 141 | $C_2$: Ala 118 | $A_2$: Ala 118 | $B_2$: Ala 118 | C: Asn 96 | $A_2$: Asn 96 |
| | B: Thr 142 | | | $C_2$: Ser 119 (h) | $A_2$: Ser 119 (h) | $B_2$: Ser 119 (h) | C: Gly 97 | $A_2$: Gly 97 |
| | B: Ile 143 | A: Ile 143 | C: Ile 143 | $C_2$: Ser 120 | $A_2$: Ser 120 (h) | $B_2$: Ser 120 (h) | C: Met 98 | $A_2$: Met 98 |
| | B: Asn 144 | A: Asn 144 | C: Asn 144 | $C_2$: Gln 121 | $A_2$: Gln 121 (h) | $B_2$: Gln 121 (h) | C: Lys 100 | $A_2$: Lys 100 |
| | | A: Phe 156 | C: Phe 156 | $C_2$: Asn 122 | $A_2$: Asn 122 | $B_2$: Asn 122 | C: Ala 101 | $A_2$: Ala 101 |
| | B: Ser 159 | A: Ser 159 | C: Ser 159 | | $A_2$: Thr 149 | $B_2$: Thr 149 (h) | | $A_2$: Tyr 102 |
| | B: Tyr 160 | A: Tyr 160 | C: Tyr 160 | | $A_2$: Pro 151 | $B_2$: Pro 151 | | $A_2$: Arg 191 |
| | B: Asn 162 | A: Asn 162 | C: Asn 162 | $C_2$: Ser 185 | $A_2$: Ser 185 | $B_2$: Ser 185 | | $A_2$: Phe 198 |
| | B: Thr 164 (h) | A: Thr 164 (h) | C: Thr 164 (h) | $C_2$: Ile 186 (h) | $A_2$: Ile 186 | $B_2$: Ile 186 | | $A_2$: Pro 201 |
| | B: Glu 165 | A: Glu 165 | C: Glu 165 | $C_2$: Ser 189 (h) | $A_2$: Ser 189 | $B_2$: Ser 189 | | $A_2$: Lys 202 |
| | | | | | $A_2$: Arg 191 | $B_2$: Arg 191 (h) | | |
| | C: Gly 62 | | | | | | | B: Ser 64 (s) |
| | C: His 103 (s) | B: His 103 (s) | A: His 103 (s) | B: Ser 120 (h) | A: Ser 120 (h) | C: Ser 120 (h) | | B: Glu 65 |
| | C: Glu 104 (s) | B: Glu 104 | A: Glu 104 (s) | B: Gln 121 | A: Gln 121 (h) | C: Gln 121 (h) | | B: Thr 66 |
| | C: Glu 165 (h) | B: Glu 165 (h) | A: Glu 165 (h) | B: Asn 122 (h) | A: Asn 122 | C: Asn 122 | $C_2$: Phe 67 | B: Phe 67 |
| | C: Trp 166 | B: Trp 166 | A: Trp 166 | B: Ser 123 (h) | A: Ser 123 | C: Ser 123 | | B: Val 68 |
| | C: His 167 | B: His 167 | A: His 167 | B: Gly 124 (h) | A: Gly 124 (h) | C: Gly 124 (h) | | B: Cys 91 |
| | C: Asp 168 | B: Asp 168 | A: Asp 168 | B: Ser 125 (h) | A: Ser 125 (h) | C: Ser 125 | $C_2$: Pro 92 | B: Pro 92 |
| | C: Ala 170 | B: Ala 170 | A: Ala 170 | | A: Tyr 128 | | $C_2$: Ala 93 | B: Ala 93 |
| | C: Glu 171 (s) | B: Glu 171 | A: Glu 171 (s) | B: Thr 142 (h) | A: Thr 142 (h) | C: Thr 142 (h) | $C_2$: Asn 96 | B: Asn 96 |
| | C: His 199 | B: His 199 | A: His 199 | B: Ile 143 | A: Ile 143 | C: Ile 143 | $C_2$: Gly 97 | B: Gly 97 |
| | C: Asn 200 (h) | B: Asn 200 (h) | A: Asn 200 (h) | | A: Asn 144 | C: Asn 144 (h) | $C_2$: Met 98 | B: Met 98 |
| | C: Pro 201 | B: Pro 201 | A: Pro 201 | B: Lys 145 (hs) | A: Lys 145 (s) | C: Lys 145 (s) | $C_2$: Lys 100 | B: Lys 100 |
| | C: Lys 202 (h) | B: Lys 202 (h) | A: Lys 202 | | | C: Phe 146 | $C_2$: Ala 101 | B: Ala 101 |
| | | | | | A: Gly 147 | C: Gly 147 | $C_2$: Tyr 102 | B: Tyr 102 |
| | | | | | A: Thr 149 | C: Thr 149 | | B: Arg 191 |
| | | | | | A: Lys 150 (h) | C: Lys 150 (h) | | B: Phe 198 |
| | | | | | A: Arg 154 | C: Arg 154 | | B: Pro 201 |
| | | | | B: Lys 179 | | | | B: Lys 202 |
| | | | | B: Asn 181 (h) | A: Asn 181 (h) | C: Asn 181 (h) | | |
| | | | | B: Gly 182 | | B: Gly 182 | | |

§ Chains and rotation axis labelled as in Fig. S3e.
† (h) and (s) indicate residues involved in interfacial hydrogen bonds and salt bridges respectively.

**Table S5. Residues at the interfaces between subunits.**

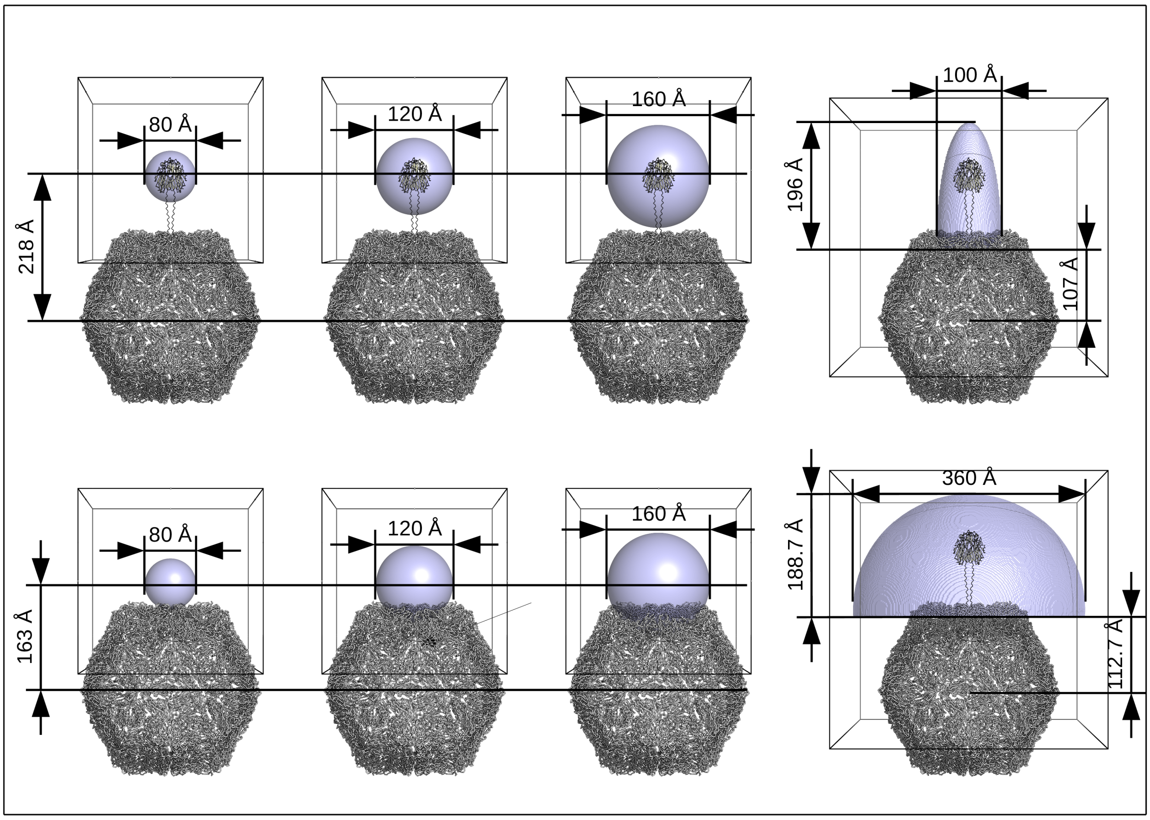


**Fig. S1. The masks that have been used for asymmetric reconstructions of the $^{N}$RTD.** The masks shown in the figure are focussed around an $I_2$ axis (vertical direction). Masks of the same shapes and sizes, aligned with the closest $Q_2$ and $Q_3$ axes have also been used. For reference, the putative farthest position of the $^{N}$RTD from the capsid is shown assuming a polyproline-like helical conformation of the CP linkers parallel to the symmetry axis. Note that the $^{N}$RTD model was never used as a reference during 3D classifications.

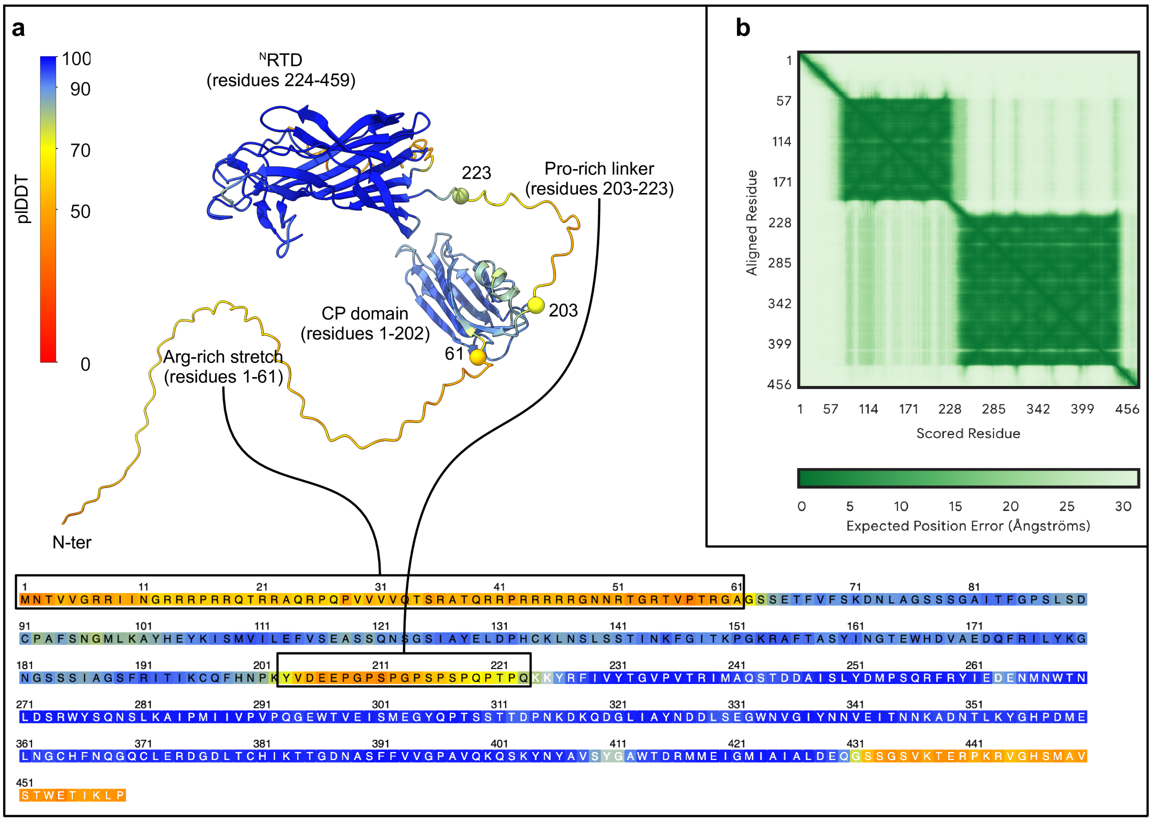


**Fig. S2. AlphaFold-3 model of the TuYV RTP* monomer.**

(**a**) Structure of the RTP* monomer predicted by AlphaFold-3. The structure and its sequence are coloured according to the confidence estimate (plDDT). The Arg-rich stretch and Pro-rich linker are zones of low-confidence structural prediction (plDDT<70). (**b**) Estimate of the error in the relative position and orientation between two tokens in the predicted structure. High values (low confidence) are found for the position of the CP jelly-roll domain relative to the $^{N}$RTD.

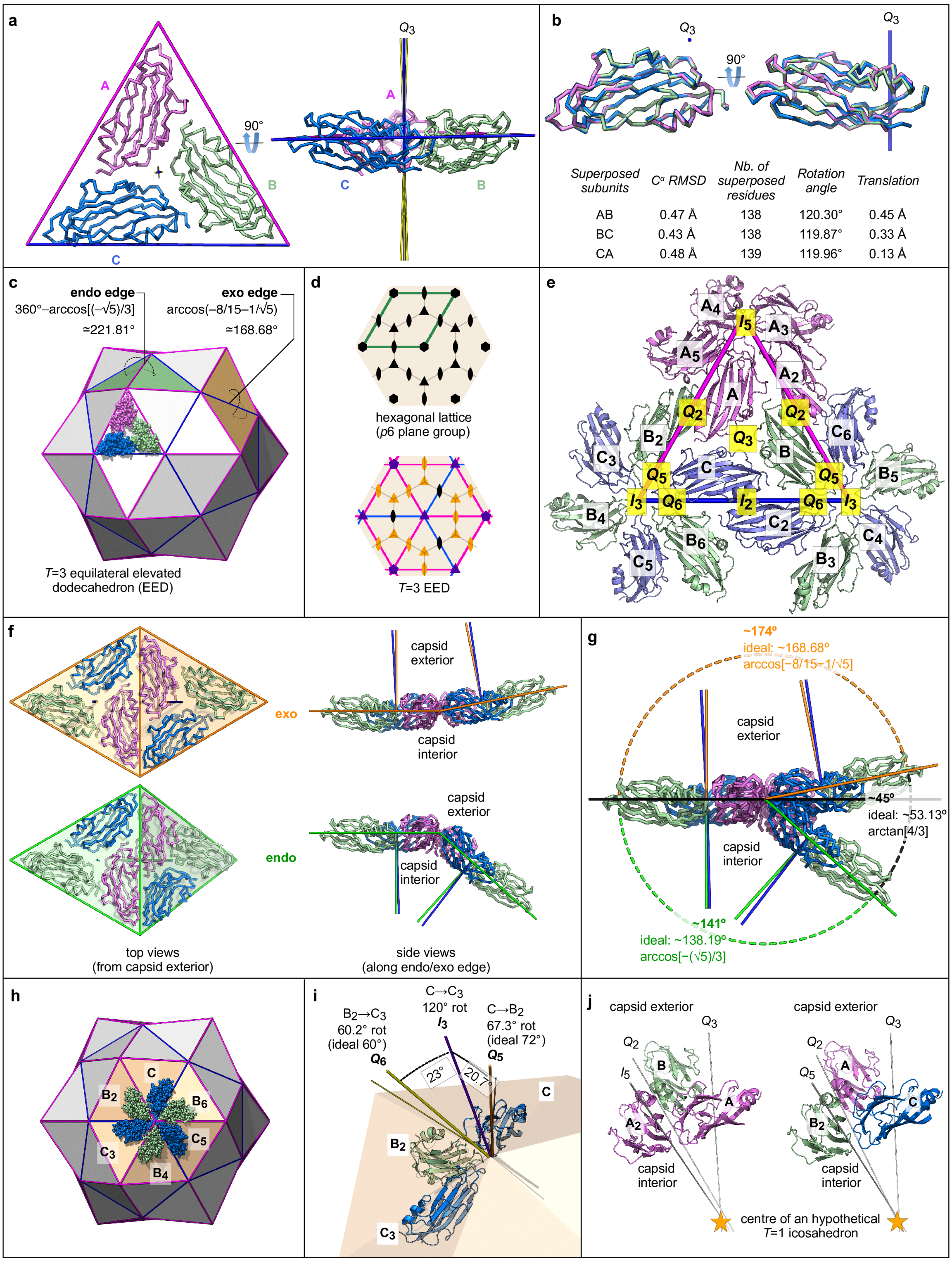


| Superposed subunits | $C^\alpha$ RMSD | Nb. of superposed residues | Rotation angle | Translation |
|---|---|---|---|---|
| AB | 0.47 Å | 138 | 120.30° | 0.45 Å |
| BC | 0.43 Å | 138 | 119.87° | 0.33 Å |
| CA | 0.48 Å | 139 | 119.96° | 0.13 Å |

### Fig. S3. Architecture of the TuYV capsid.

#### Fig. S3 (caption)

(**a**) Top view from the capsid exterior (left) and side view (right) of one asymmetric unit of the icosahedral assembly. The three subunits (chains A, B and C) are pairwise related by almost exact threefold axes (yellow rods). An ideal threefold axis (calculated by a least-squares fitting procedure) is indicated by a blue vertical rod and labelled $Q_3$. (**b**) Two views of the three subunits superposed onto each other and oriented as chain C in (a). The ideal threefold axis ($Q_3$) is shown in blue. The results of pairwise superpositions of the asymmetric unit chains are reported. (**c**) Representation of the equilateral elevated dodecahedron (EED) used for describing the TuYV capsid. The three subunits in one asymmetric unit are also shown. Two pairs of triangular faces, one in *endo* and the other in *exo* configuration, are highlighted in green and orange respectively. *Endo* and *exo* edges are coloured magenta and blue respectively. (**d**) Symmetry diagram of a virtual hexagonal lattice (top) and the resulting EED lattice (bottom) upon folding along the *endo* (magenta) and *exo* (blue) edges. The colours of the symmetry symbols in the bottom picture indicate how the nature of the symmetry axes changes upon folding: black means transformation into an icosahedral axis of the same order as in the hexagonal lattice, violet means transformation into an icosahedral axis of a different order, and orange means transformation into a local symmetry axis. (**e**) Arrangement of subunits around one asymmetric unit viewed from the capsid exterior. Subunits related by an icosahedral rotation are coloured the same and labelled with the same letter, while quasi equivalent subunits not related by icosahedral rotations are coloured differently and labelled with different letters. The approximate positions of icosahedral *n*-fold ($I_n$, $n = 2, 3, 5$) and local-symmetry quasi *n*-fold ($Q_n$, $n = 2, 3, 5, 6$) axes are indicated by the yellow labels. The triangle delimits one icosahedral asymmetric unit. *Endo* edges in magenta, *exo* edge in blue. (**f**) Two pairs of asymmetric units forming an *exo* (top) or *endo* edge (bottom). On the left: the triangular faces of and ideal EED are also shown. The leftmost faces are orthogonal to the view direction. On the right: the directions perpendicular to the faces of an ideal EED are indicated by the orange or green rods, while the actual local threefold axes of TuYV are indicated by the blue rods. (**g**) An *endo* and an *exo* pairs of asymmetric units represented superposed. The view direction is along the axis of the virtual hinge movement that would transforms an *endo* angle into an *exo* angle. (**h**) Hexameric arrangement of TuYV subunits around an $I_3$ axis of the ideal EED. Only subunits coloured the same are related by the icosahedral threefold symmetry. EED edges coloured as in (c). (**i**) A portion of a hexamer (three adjacent subunits) and the rotation axes relating pairs of these subunits. The thicker rods, labelled $Q_6$, $I_3$ and $Q_5$, indicate the actual axis directions in the TuYV capsid. The two thinner rods close to the $Q_6$ and $Q_5$ axes represent the directions of the corresponding axes in an ideal EED. For reference, the six EED faces underlying a complete hexamer are partly visible. (**j**) Two quasi-equivalent triplets of subunits, showing that the quasi-symmetry axes relating their chains would be compatible with those of a $T$=1 icosahedral assembly.

**a**

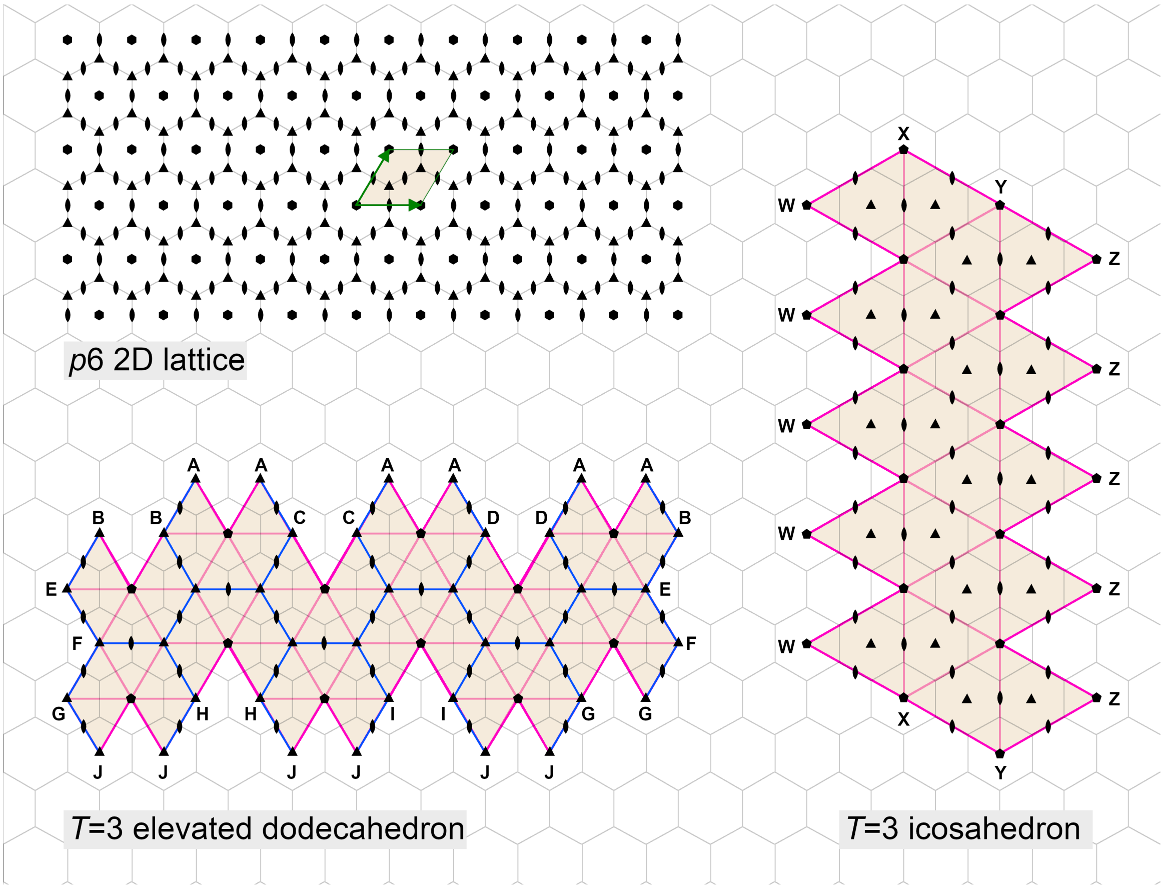


**b**

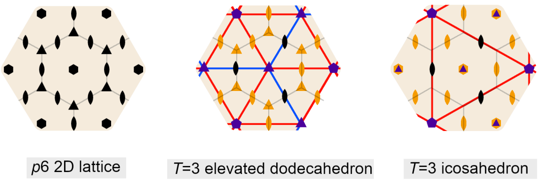


**Fig. S4. Folding of a hexagonal lattice into either an icosahedron or an elevated dodecahedron.**

(**a**) Folding of a 2D hexagonal lattice (plane space group *p*6) into a *T*=3 icosahedron or a *T*=3 equilateral elevated dodecahedron (EED). Folding along the blue (*exo* angles) and magenta edges (*endo* angles) generates the two polyhedra. A unit cell of the 2D lattice is highlighted (green edges). The position of the symmetry axes of the plane lattice and those of the resulting polyhedra are shown with black symbols. The lattice points that become identical upon folding are labelled with the same letter. (**b**) Transformation of the symmetry axes upon folding. Axes of the plane lattice that become icosahedral symmetry axes without changing their order are coloured black in the middle and right pictures. Axes that become icosahedral symmetry axes of a different order are coloured purple. Axes that become local symmetry axes are coloured orange.

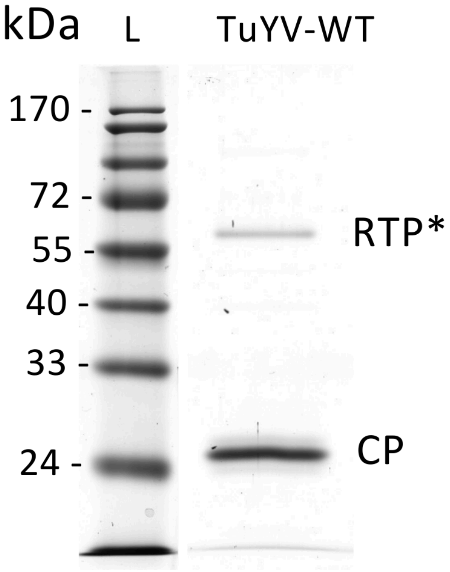


**Fig. S5. SDS-PAGE analysis and colloidal Coomassie blue staining of 2 µg of TuYV-WT viral particles purified from *M. perfoliata* and denaturated in Laemmli buffer.**
L: Ladder.

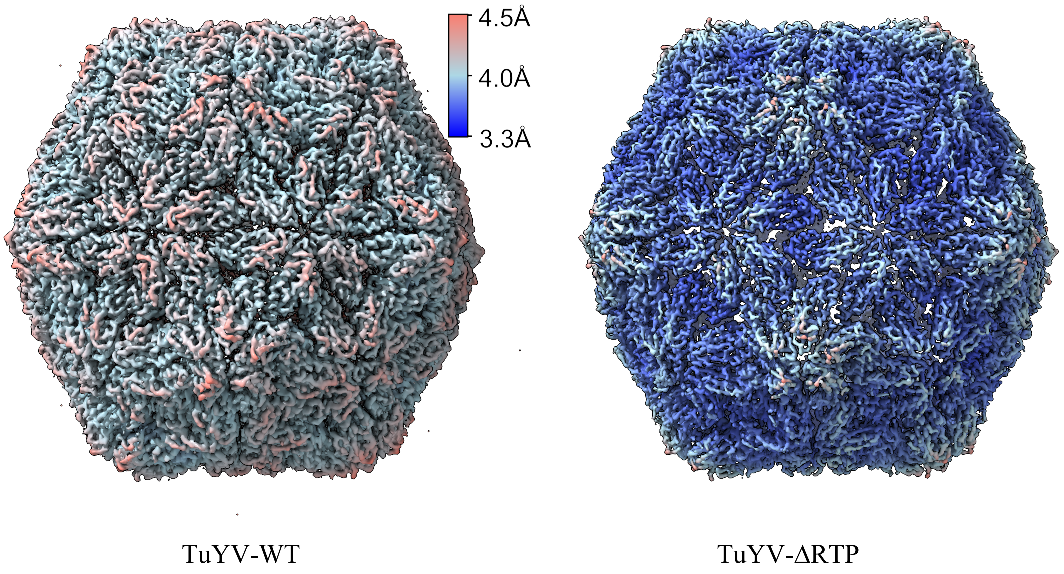


**Fig. S6. Local resolution of the cryo-EM reconstructions (sharpened maps).**

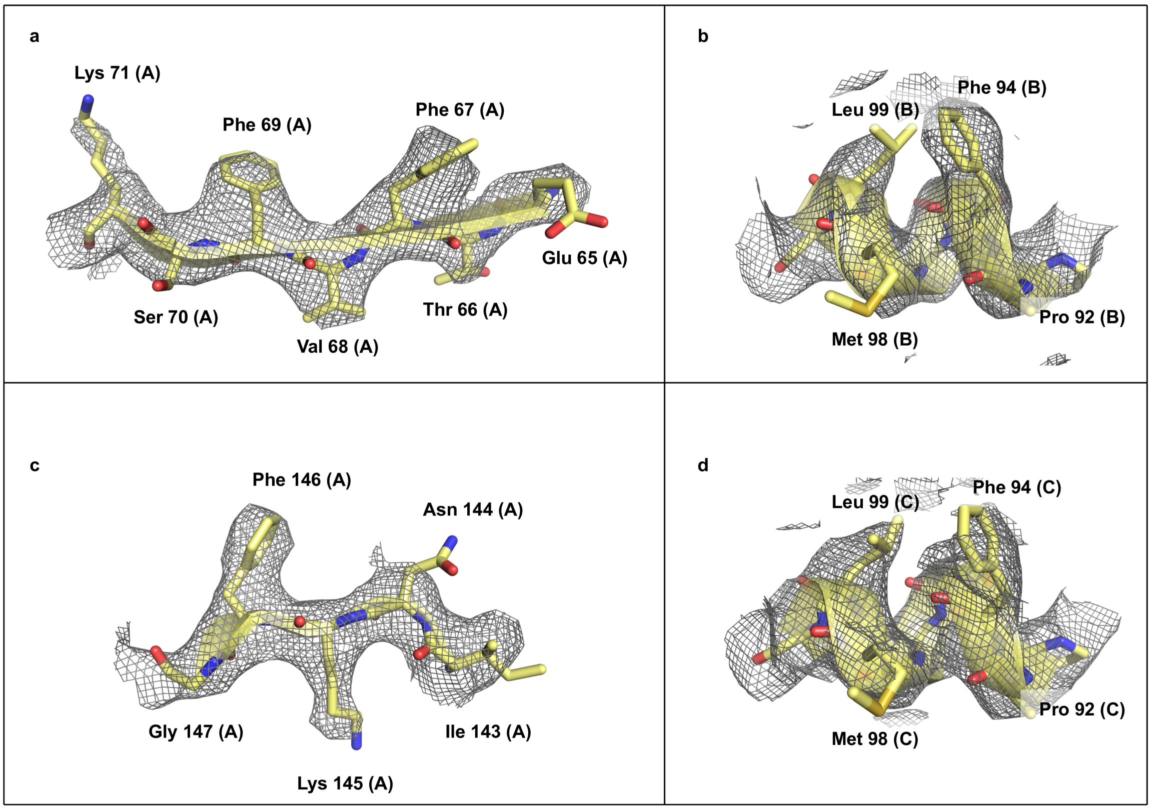


**Fig. S7. Some details of the TuYV-ΔRTP cryo-EM map and the corresponding refined atomic model.**

Regions (a) and (b) are involved in $Q_2$ subunint-subunit interfaces, region (c) is involved in $I_5$ interfaces, and region (d) is involved in $I_2$ interfaces.

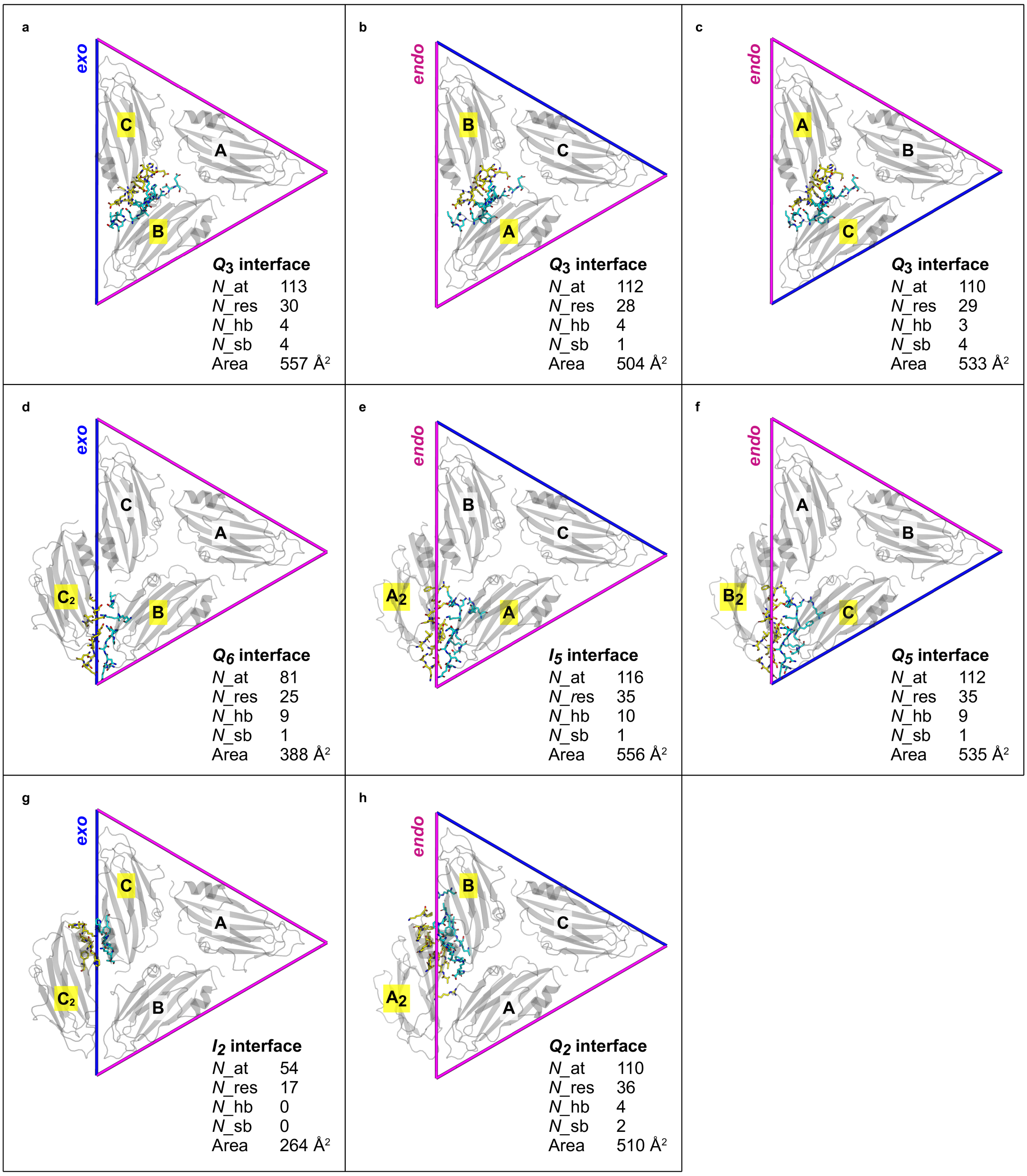


**Fig. S8. The eight non-equivalent subunit interfaces.**

In each panel a triangular asymmetric unit of the capsid (chains A, B and C) and, eventually, an additional neighbouring chain are depicted. Subunits are labelled as in Fig. S3. The *endo* and *exo* edges delimiting the asymmetric unit are coloured magenta and blue respectively. The viewpoints in panels b+e+h and and c+f differ from the viewpoint in panels a+d+g by a clockwise in-plane rotation of 120° and 240° respectively. In each panel, the interface between two subunits (yellow labels) is highlighted. Interfacial residues are represented in coloured stick mode. N_at: number of interfacial atoms. N_res: number of interfacial residues. N_hb: number of interfacial hydrogen bonds. N_sb: number of interfacial salt bridges.

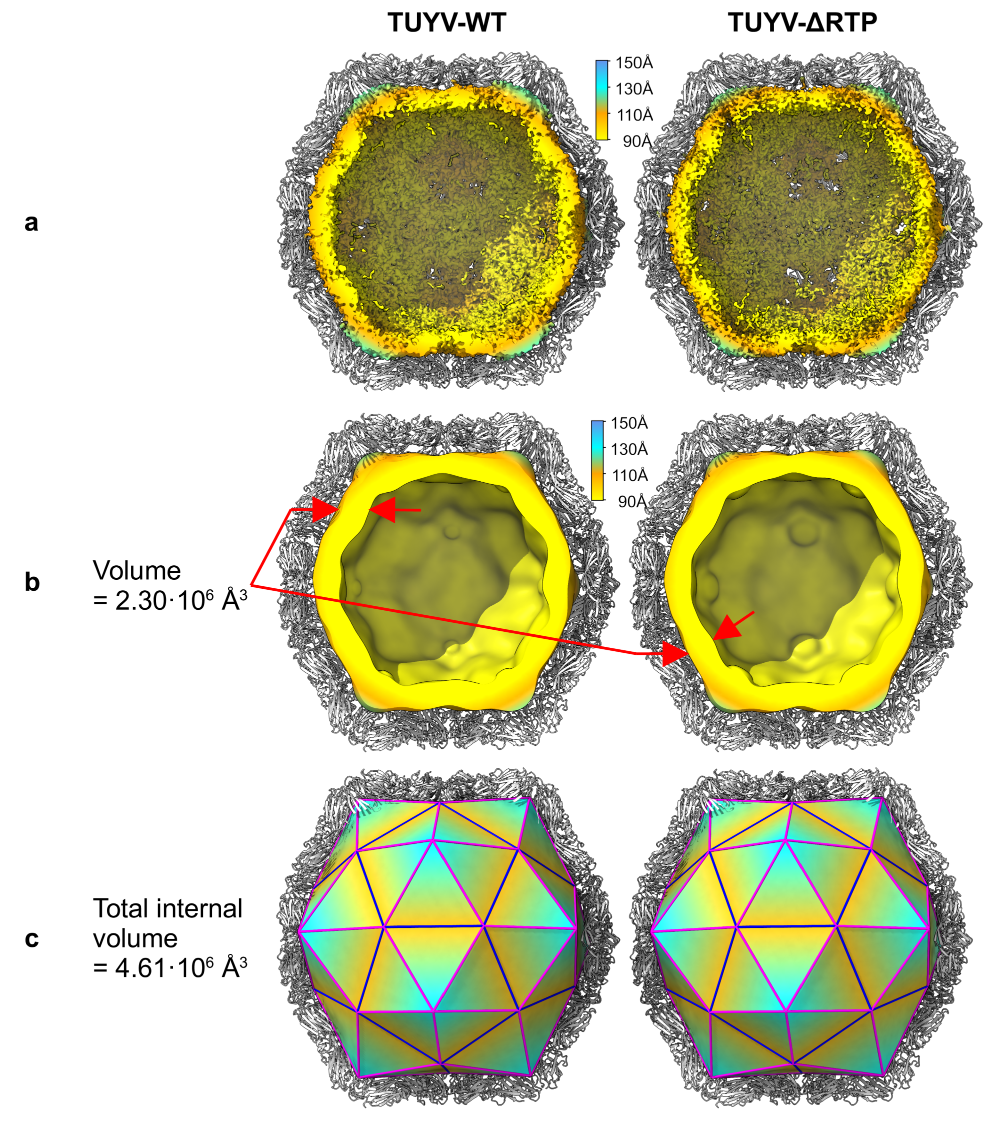


d

| | Sequence-based volume estimations | |
|---|---|---|
| Genome | 1 717 293 Å³ (5 641 nucleotides) | 1 323 664 Å³ (4 348 nucleotides) |
| CP N-ter Arg-rich stretches | 543 552 Å³ (60 x 63 amino acids) | 543 552 Å³ (60 x 63 amino acids) |
| **Total** | **2 260 845 Å³** | **1 867 216 Å³** |

**Fig. S9. Estimation of the residual density volumes at the capsid interior.**
(**a**) The residual density (in colours) after masking out the contribution of the CP shell (white cartoon model). The density is coloured according to the distance from the particle centre. (**b**) Low-pass filtered residual density. The contour level has been chosen so that the enclosed volume encompasses the predicted volume of the genomic RNA plus the volume of the unmodelled Arg-rich N-terminal stretches of the CP. (**c**) The volume of an ideal EED inscribed within the capsid shell. (**d**) Sequence-based estimation of the volume occupied by the unmodelled atoms at the interior of the capsid.
